# Whole genome sequencing reveals fine-scale climate associated adaptive divergence near the range limits of a temperate reef fish

**DOI:** 10.1101/2022.11.28.517507

**Authors:** Cameron M. Nugent, Tony Kess, Matthew K. Brachmann, Barbara L. Langille, Steven J. Duffy, Sarah J. Lehnert, Brendan F. Wringe, Paul Bentzen, Ian R. Bradbury

**Affiliations:** Fisheries and Oceans Canada, Northwest Atlantic Fisheries Centre, St. John’s, Newfoundland, Canada; Fisheries and Oceans Canada, Bedford Institute of Oceanography, Dartmouth, Nova Scotia, Canada; Dalhousie University, Department of Biology, Halifax, Nova Scotia, Canada

## Abstract

Adaptation to ocean climate is increasingly recognized as an important driver of diversity in marine species despite the lack of physical barriers to dispersal and the presence of pelagic stages in many taxa. A robust understanding of the genomic and ecological processes involved in structuring populations is lacking for most marine species, often hindering management and conservation action. Cunner (*Tautogolabrus adspersus*), is a temperate reef fish that displays both pelagic early life history stages and strong site-associated homing as adults; the species is also presently of interest for use as a cleaner fish in salmonid aquaculture in Atlantic Canada. Here we produce a chromosome-level genome assembly for cunner and characterize spatial population structure throughout Atlantic Canada using whole genome resequencing. The genome assembly spanned 0.72 Gbp and resolved 24 chromosomes; whole genome resequencing of 803 individuals from 20 locations spanning from Newfoundland to New Jersey identified approximately 11 million genetic variants. Principal component analysis revealed four distinct regional groups in Atlantic Canada, including three near the range edge in Newfoundland. Pairwise *F*_ST_ and selection scans revealed consistent signals of differentiation and selection at discrete genomic regions including adjacent peaks on chromosome 10 recurring across multiple pairwise comparisons (*i.e*., *F*_ST_ 0.5-0.75). Redundancy analysis suggested significant association of environmental variables related to benthic temperature and oxygen range with genomic structure, again highlighting the previously identified region on chromosome 10. Our results suggest that climate associated adaptation in this temperate reef fish drives regional diversity despite high early life history dispersal potential.

## INTRODUCTION

Genetic structuring of marine populations and adaptive differentiation has been shown to evolve despite high dispersal potential as a result of the interplay of environment, life history, and other factors influencing survival. Many marine taxa have life histories that feature pelagic egg and/or larval stages, enabling current-mediated dispersal across large distances which is thought to contribute to the low levels of genetic structuring often observed in marine fish (Levin 1996; Waples 1998; Juanes 2007; Knutsen *et al*. 2022). However, low levels of genetic structuring are not universal in marine species (Schmidt *et al*. 2008) and environmental gradients have been repeatedly associated with regional genetic structure, suggesting that adaptive evolutionary change can occur in response to local environmental differences (De Faveri *et al*. 2013; Duranton *et al*. 2018; Jahnke *et al*. 2018; Stanley *et al*. 2018; Knutsen *et al*. 2022; Pratt *et al*. 2022). This is especially true in harsh environments such as range edges, where an individual’s ability to tolerate harsh conditions is essential to survival (Schmidt *et al*. 2008; Blank *et al*. 2013; Sylvester *et al*. 2018). Ultimately, understanding the distribution of adaptive diversity in marine species is essential to successful management and conservation of marine resources particularly those threatened by anthropogenic impacts including exploitation and climate change.

Population genomics-based examinations of diversity in marine species are revealing the geographic and genomic distribution of differentiation and potential adaptive diversity. Studies using large SNP panels (Sylvester *et al*. 2018; Watson *et al*. 2021), reduced representation sequencing (Pratt *et al*. 2022; Sønstebø *et al*. 2022), and whole genome resequencing approaches (Han *et al*. 2020; Knutsen *et al*. 2022) have with increasing levels of detail revealed significant climate or environmental associations that localize to discrete genomic regions. For example, recent work combining pooled whole genome sequencing and double-digest sequencing methods, characterized the spatial genetic structure of broad-nosed pipefish (*Syngnathus typhle*). Additionally, comparison to other species with different life-history characteristics demonstrated the roles of fragmented habitat and an absence of a pelagic larval stage in promoting genetic structure (Knutsen *et al*. 2022). Another recent study of two related redfish species (*Sebastes mentella* and *Sebastes fasciatus*) utilized SNPs identified through reduced representation sequencing to assess population genomic structure and reveal its association with spatial distribution (Benestan *et al*. 2021). Research has shown that key regions of the genome harbouring putative adaptive diversity are often influenced by the interplay of genomic structural variation, environment associated selection, recombination, and drift (Moore *et al*. 2014; Sylvester *et al*. 2018; Han *et al*. 2020; Tigano *et al*. 2021; Watson *et al*. 2021; Knutsen *et al*. 2022; Pratt *et al*. 2022). Improving clarity of these potential genomic islands of adaptive diversity using high resolution genome sequencing methods can directly inform efforts to manage exploited marine species and help conserve at risk populations (Funk *et al*. 2012).

Cunner (*Tautogolabrus adspersus*) are a species of Labridae whose range covers a wide gradient of ecological variation from Chesapeake Bay, USA to the northern edge of Newfoundland, Canada (Moran *et al*. 2019). Like other temperate wrasses, cunner are physiologically adapted to survive at colder temperatures, annually entering a prolonged state of physiological torpor when water temperatures fall below 5°C, which in the northern portion of its range may last 6 months (Bradbury *et al*. 1995; Moran *et al*. 2019). Cunner display pelagic egg and larval stages, both of which occur during summer months in Atlantic Canada (Bradbury *et al*. 2003). Adults have been shown to occupy small ranges and accurately home following displacement (Green 1975). As such, there is potential for dispersal in this species, but the genetic structure of cunner has not been characterized in Atlantic Canada. Cunner are presently being considered as a candidate species for use as a cleaner fish within Atlantic salmon net pen aquaculture in Atlantic Canada (Kelly Cove Salmon Ltd. 2012; Chen 2020) where sea lice continue to hamper industry production and growth. However, cunner escapees from net pens could influence the fitness and demography of wild populations (*i.e*, Faust *et al*. 2018; Faust *et al*. 2020) and recent evidence suggests escaped cleaner fish can interbreed with wild conspecifics (*e.g*. Wringe *et al*. 2018). Understanding the potential ecological and genetic impacts of cunner aquaculture escapees on wild populations is an important component to consider as the use of the species in salmon aquaculture production is further developed (Naylor *et al*. 2005; Karlsson *et al*. 2016; Blanco Gonzalez & de Boer 2017).

Our main objective was to use whole genome re-sequencing (WGS) to evaluate the geographic and genomic scale of differentiation for cunner in the Northwest Atlantic. The specific goals were to: 1) develop genomic resources for cunner through the creation and quality assessment of the first chromosome-level genome assembly in this species; 2) use this reference genome in conjunction with individual WGS data to explore population level differentiation present in the Northwest Atlantic and its distribution both in space and across the genome; 3) and finally explore the potential drivers of differentiation in this species using environmental association analysis. This research extends previous work quantifying genomic-environmental associations in other marine taxa in the region (Stanley *et al*. 2018; Layton *et al*. 2021; Kess *et al*. 2021; Watson *et al*. 2022) and provides one of the first whole genome evaluations of climate associated adaptation in a marine species within Atlantic Canada. The goal was to characterize genomic diversity and population structure across the sampled range and thereby understand the evolutionary history of the species at the northern extreme of their range. Through the wide sampling range and fine scale resolution provided by the WGS methods employed, we aimed to understand the forces shaping genomic diversity in cunner, a marine species with high egg and larval dispersal potential as well as local site fidelity as adults.

## MATERIALS AND METHODS

### Sample and tissue collection

Between July 8^th^ and October 19^th^, 2019, sampling was conducted at 19 locations throughout Atlantic Canada (Table 1; Figure 1) using a combination of angling and traps, with a maximum of 100 individuals collected per location. In total, 1379 fin clip samples were obtained from the sampling locations and a representative subset were selected for DNA extraction and sequencing. An additional 54 individuals from Tuckerton, New Jersey (obtained from pelagic larvae collections in 2012) were included in the set of sequenced samples.

**Table 1.**
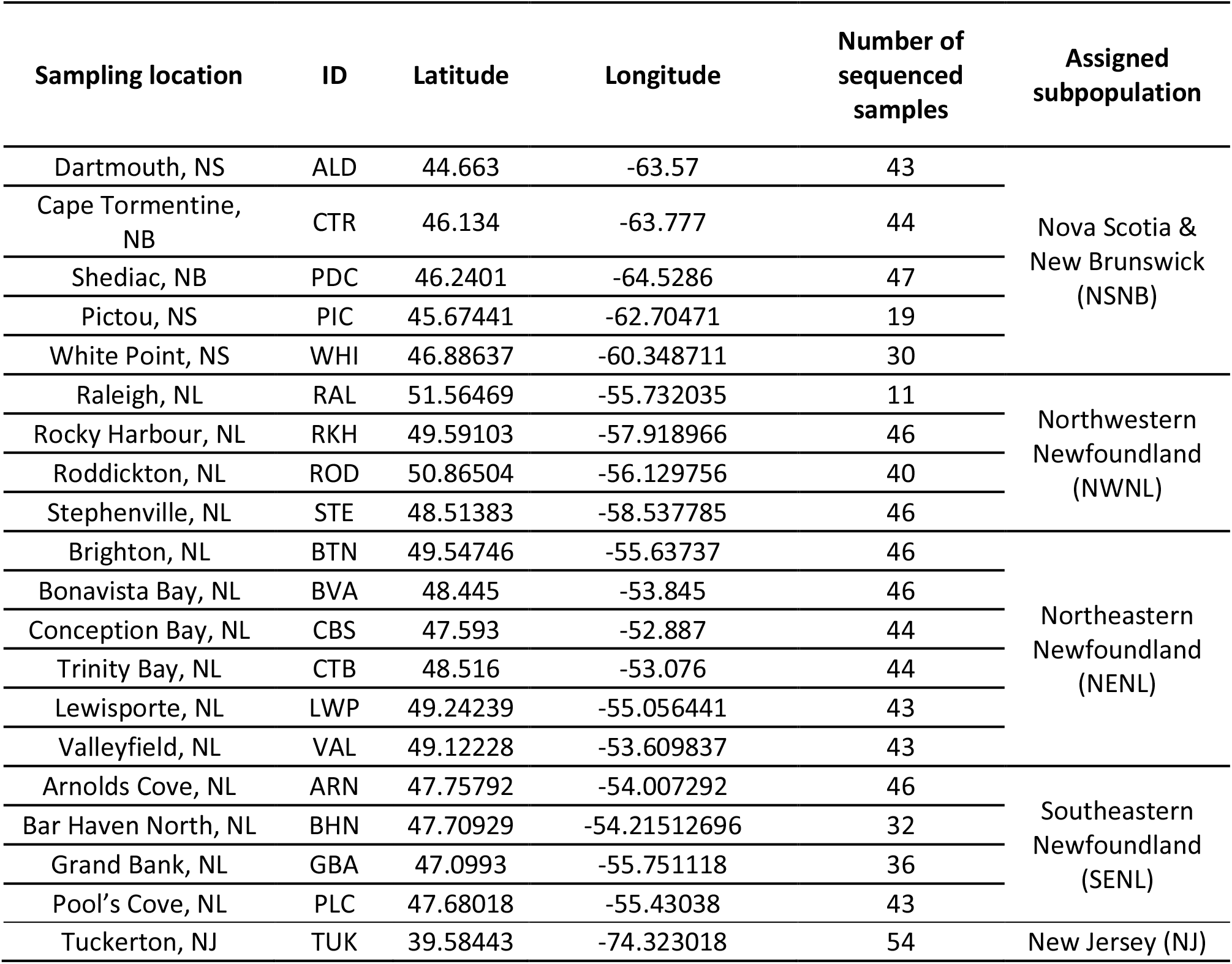
Cunner sampling locations and sample sizes used in the analyses of population structure and genomic diversity. The ALD population initially consisted of 45 sequenced samples, but analysis of the genetic information was suggestive of the two samples being duplicates of each other. Conservatively, these samples were therefore both removed and only 43 samples were used in subsequent analyses. The Assigned subpopulation indicates the predominant *k*-means cluster assignments of individuals from each location (which averaged 96.5% intra-location concordance), deviations from the majority location assignment can be observed in Table S2)

**Figure 1.**
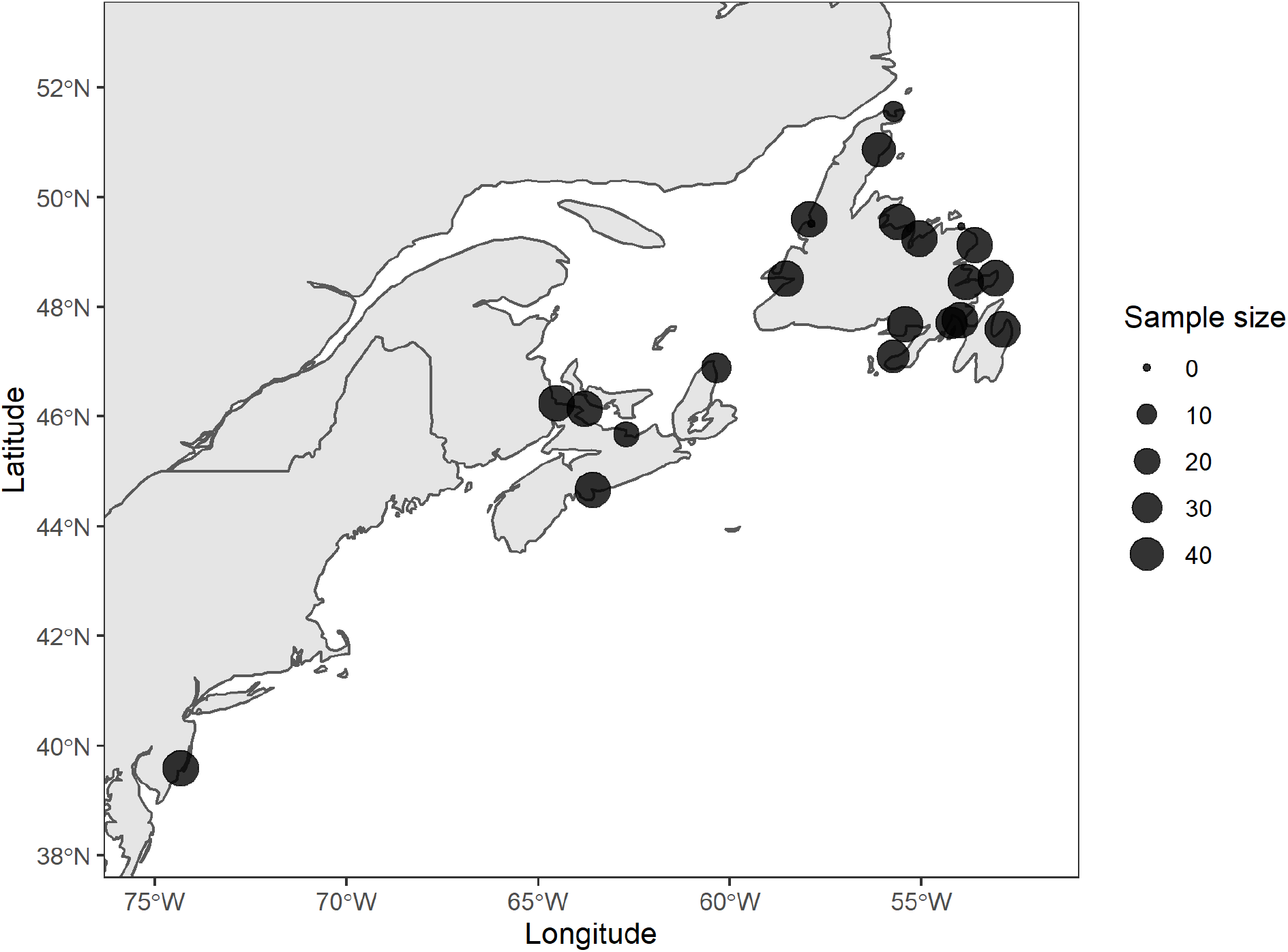
Map of cunner sampling locations. The size of the points is reflective of the number of samples from each location that were sequenced and used in population genomic analyses. For more information on sampling locations see Table 1.

### Reference genome assembly and quality assessment

#### Genome sequencing and assembly

Muscle tissue from a single male adult cunner collected from Alderney Landing in Dartmouth, Nova Scotia, Canada on August 12, 2019, was used for high-quality reference assembly generation as part of the Vertebrate Genome Project (VGP). Library preparation, sequencing, and assembly were performed following the VGP standard assembly pipeline version 1.6 (see Rhie *et al*. 2021 for assembly methods and software details). Sequence information for the reference individual was generated through a combination of PacBio long reads (103.8x coverage), 10x Genomics Illumina reads (88.9x), BioNano optical maps (744.5x), and Arima Hi-C Illumina read pairs (64.9x) and then assembled through the VGP pipeline which is composed of an initial assembly step, followed by scaffolding and final polishing (Rhie *et al*. 2021).

#### Genome quality assessment

To assess the completeness and quality of the assembled genome, a Benchmarking Universal Single Copy Orthologs (*BUSCO*, version 5.2.2.) analysis was conducted. The “actinopterygii_odb10” lineage dataset, which contains 3640 genes derived from 26 species, was used for BUSCO analysis. The following parameters were used: *“-m genome -l actinopterygii_odb10*”. BUSCO was run in genome mode and the gene predictor used was *metaeuk* (Levy *et al*. 2020).

To characterize the distribution of repetitive elements throughout the genome, a *de novo* repeat library was constructed by querying the genome using *RepeatModeler* (v2.0.2), along with the dependencies *RepeatScout* (v1.0.5), *rmblast* (v2.10.0+), and *RECON* (v1.0.8). Library construction was conducted using default parameters over five analysis rounds, along with the LTR discovery pipeline option (*-LTRStruct*). *RepeatMasker* (v4.1.1) was used along with the repeat library to predict and annotate repetitive elements present within the genome (Smit *et al*. 2013).

#### Comparative genomics with other wrasses

Whole genome alignment of the cunner reference genome to two closely related wrasse species, humphead wrasse (*Cheilinus undulatus*; Liu *et al*. 2021; accession number: GCF_018320785.1) and New Zealand spotty wrasse (*Notolabrus celidotus*; Rhie *et al*. 2021; accession number: GCF_009762535.1) was conducted to investigate the quality of the cunner genome assembly, identify homologous chromosomes within other species, and search for evidence of differences in genomic architecture.

Megablast was used to align the 24 cunner chromosomes (query) to the other wrasse species (subject) with the following paramteres: *‘-evalue 0.0001 -max_target_seqs 3 -max_hsps 20000 -outfmt 6 - num_threads 32 -word_size 40 -perc_identity 80’*. Following megablast, outputs were filtered using a custom Python script adapted from Christensen *et al*. 2018 (from source manuscript, Supplementary script: “Compare_Genome_2_Other_Genome_blastfmt6_ver1.0.py” run with the parameters: *‘-minl 0.01 -minal 5000’*). The results were visualized in the R programming language and homology blocks tabulated to identify putative homologies. For completeness, the process was repeated aligning the New Zealand spotty wrasse (query) to the humphead wrasse genome (subject).

#### Whole genome resequencing

We conducted whole genome resequencing (WGS) for cunner collected from 20 sampling locations (Table 1). Fin clips from individuals were preserved in 95% ethanol and subsequent DNA extraction was conducted using DNeasy 96 Blood and Tissue kits (Qiagen) according to manufacturer’s protocols. The quality of extracted DNA was visualized using 1% agarose gel electrophoresis and quantified using Quant-iT PicoGreen ds-DNA Assay kits (Thermofisher) and a fluorescent plate reader. DNA samples were normalized to 15ng / μl. Paired-end whole genome sequencing was conducted on nine lanes of an Illumina NovaSeq6000 S4 at The Genome Quebec Centre d’Expertise et de Services. Manual inspection of resulting file sizes was conducted and samples with forward or reverse read files less than 10% of the average sequence file size for the whole set were considered problematic and removed. The remaining 805 samples were processed using a custom bioinformatics pipeline based on the methods initially developed for Atlantic salmon WGS processing (https://github.com/TonyKess/seaage_GWAS). All scripts used in cunner WGS processing are publicly available on GitHub (https://github.com/CNuge/lowdepth-2-snps). The pipeline was deployed in a batch fashion on the Compute Canada Graham computing cluster, using the *slurm* scheduler (Yoo *et al*. 2003) and GNU parallel to facilitate parallelized computation (Tang 2018).

Paired end sequence files were processed using *cutadapt* (version 3.4, Martin 2011) to remove the leading 15 base pairs, known adapter sequences, and base pairs with quality (Q scores) below 10. Any reads less than 40bp in length following trimming were removed entirely. For each sample, *bwa* (version 0.7.17, Li & Durbin 2009) was used to conduct a Burrows-Wheeler alignment of the paired end sequence reads against the cunner reference genome assembly (GCA_020745685.1). Resulting bam files were sorted with *samtoools* (version 1.13, Danecek *et al*. 2021) and deduplicated using *picard*’s MarkDuplicates function (version 2.26.3, Broad Institute 2019). Using *gatk* (version 3.8, McKenna *et al*. 2010) indel realignment was conducted via the RealignerTargetCreator and IndelRealigner functions. The realigned bam files were input to *angsd* (version 0.933, Korneliussen *et al*. 2014) for genotyping, genotype likelihood estimation, and data filtering through the inclusion of the following parameters: *-minMaf 0.01, -minMapQ 30, -minQ* 20, as well as *-minInd* and *-setMinDepth* both set to 80% of the total individual count. The genotyping and genotype likelihood estimation was conducted on a per chromosome basis.

### Characterizing genomic variation within and between populations

#### Identifying population substructure with Principal Component Analysis

The program *PCAngsd* (version 1.02, Meisner & Albrechtsen 2018) was used to quantify population structure. This was done twice using matching methods and different data sets: first the complete set of 805 samples was considered to assess structure across the entirety of the sampled range, second the 751 samples from the Atlantic Canada sampling locations were analyzed to allow for finer scale resolution of population structure within this region of interest. For each analysis, the by-chromosome beagle files containing genotype likelihoods for the samples were merged into a single file and *PCAngsd* was used to estimate the covariance matrix of the genotype likelihoods. The resulting matrix was imported into the R programming environment and principal components (PCs) were obtained through calculation of eigenvalues using the R function *eigen*. The percent variance explained by each principal component was calculated and used to determine the optimal number of PCs for subsequent analyses.

Population structure was determined through the *k*-means clustering of individual’s values for PCs 1 and 2. Clustering was conducted iteratively, for values of *k* from 1 to 20 and for each *k* the within cluster sum of squares (*wss*) was calculated. A scree plot was generated and used to infer the optimal value of *k* by identifying the inflection point in the *wss* trendline. Final *k*-means clustering with the optimal *k* was conduced and individuals were assigned to putative subpopulations based on their assigned cluster. This information was visualized and retained for subsequent analysis of the genetic differentiation between identified subpopulations.

#### *Assessing genomic differentiation with pairwise F*_*ST*_

To identify genomic regions showing differentiation among populations we conducted pairwise comparisons of the distinct Atlantic Canada populations identified in the clustering of PCs of genetic variation and the additional population of New Jersey samples. For each pairwise combination of populations, we used *vcftools* (version 0.1.16, Danecek *et al*. 2011) to estimate Weir & Cockherham’s *F*_ST_ (1984) on a per locus basis and the outputs were loaded into R and visualized using *ggman* (https://github.com/drveera/ggman) to identify peaks in *F*_ST_. The number and distribution of loci exhibiting *F*_ST_ values > 0.15 were quantified. Along with visual inspection of the Manhattan plots, we quantified the number of *F*_ST_ peaks using a custom Python script and examined consistency across the Atlantic Canada comparisons on a SNP-by-SNP basis. An *F*_ST_ peak was defined as: five or more loci with *F*_ST_ greater than 0.15 and no more than 100 Kb separating adjacent loci. The location of these peaks across pairwise comparisons was examined to determine their relationship to the geographic distribution of the populations.

We conducted an additional PCA of the samples using only the set of markers with *F*_ST_ > 0.15 found within the identified *F*_ST_ peaks. The relevant individuals and markers were subset from the complete beagle file, and the analysis was conducted using *PCAngsd* with the same methods described for the previous PCA. As a control comparison, we repeated this analysis using a random subset (roughly equivalent to the number of outlier loci) of 8,000 markers not found within the *F*_ST_ peaks.

#### Selection scan

To search for evidence of polygenic signals of selection between the identified cunner subpopulations, we estimated XP-nSL on a per locus basis for all pairwise population comparisons (Szpiech *et al*. 2021). XP-nSL is an extension of the single-population haplotyped-based statistic nS_L_ (number of segregating sites by length) (Ferrer-Admetlla *et al*. 2014). The XP-nSL statistic compares a query and reference population to measure evidence of selection based on elevated identity by state among haplotypes. To calculate XP-nSL, we filtered full per-chromosome vcf files using *vcftools* (version 0.1.16, Danecek *et al*. 2011) with the parameters: *--maf 0.01, --min-meanDP 5, --max-missing 0.8*. We then used *vcftools* to split the filtered files by population based on PCA cluster assignments. Missing genotypes in the per population, per chromosome vcf files were then imputed using *beagle* (version 4.1, Browning & Browning 2016) and XP-nSL values were calculated for all pairwise comparisons using the program *selscan* with the option *--unphased* (version 3.0, Szpiech & Hernandez 2014). The *selscan* function *norm* was used to calculate summary statistics on a 100 Kb window basis, determining the maximum XP-nSL score per window, along with the 1% quantile for the greatest proportion of XP-nSL scores greater than or less than 2, which are respectively indicative of selective sweeps in query and reference populations of a given pairwise comparison.

#### Linkage Disequilibrium

To search for evidence of reduced recombination and putative inversions we conducted windowed analysis of linkage disequilibrium (LD) for each of the four identified Atlantic populations. For each population, *vcftools* was used on a per chromosome basis to calculate 10 Kb window-limited LD scores (using the parameters: *--geno-r2 --ld-windowed-bp 10000*). A custom Python script (Supplementary File 1) was then used to summarize the LD data on a windowed basis, calculating the mean, median, min, and max r^2^ values of all pairwise comparisons involving query loci falling within a given 100Kbp window.

The data were visually examined in R and the summary statistics were collated with the pairwise *F*_ST_ and XP-nSL scores.

#### Association of genetic and environmental variation

We aimed to describe any existing relationships between genetic variation and environmental variation across the Atlantic Canada sampling sites. The R package *sdmpredictors* (Bosch *et al*. 2017) was used to extract a series of 49 marine environment variables for the sampling locations (Supplementary File 2); the variables were derived from Bio-ORACLE (https://bio-oracle.org/) and WorldClim (https://www.worldclim.org/) data sets. These variables were analyzed and found to display numerous significant correlations; to reduce dimensionality and account for the non-independence of the environmental variables, we conducted a PCA using the R package FactoMineR (Lê *et al*. 2008). Along with the principal components (PCs) of environmental variation, the correlations between raw input variables and the principal axes of variation were obtained to understand the primary contributors to the environmental differences between locations.

Three sets of correlations were analyzed. First, the correlation of environmental and genetic variation on a per individual basis was conducted. This was conducted in R using the *lm* function, with PC1 of the Atlantic Canada PCA of the genetic variants from within the *F*_ST_ peaks used as the response variable, and PC1 of environmental variation as the predictor variable. Linear, quadratic, and cubic models for the environmental PCs were fit and compared. Secondly, this was repeated using the PC1 from the PCA of the complete set of markers in the Atlantic Canada individuals as the response variable. Third, the environmental PCs were associated with genetic variation on a per location (as opposed to per-individual) basis. To do this, minor allele frequencies (MAF) for individuals from each location were obtained using *vcftools*. A random 1% subsample of all MAF values was taken and run through a PCA using *FactoMineR*. In the same manner as the per individual analyses, the correlation of per location PCs of MAF values (a measure of intra-population genetic composition) and environmental PCs were analyzed and visualized in R.

A series of redundancy analyses were conducted as a complimentary assessment of the relationship of environmental and genomic variation (Capblancq & Forester 2021). To assess the relationship on a per-individual basis, genotype probabilities for loci within the *F*_ST_ outlier peaks were imported into R to create the response variables for the redundancy analysis (RDA). To create a set of uncorrelated predictor variables, a series of 30 marine environment variables were analyzed and a set of 6 uncorrelated variables (r^2^ < 0.7) were selected (Table S3). The environment variables were associated with genotype information based on the location of sample origin and the RDA was run using the R package *vegan* (Dixon 2003).

Results were visualized and per-loci weightings of the first two RDA axes were inspected, with scores greater than three standard deviations from the mean being defined as outliers and indicating a marker was significantly associated with the given RDA axis.

To disentangle the effects of location and population connectivity from environmental effects on genetics, the per-individual RDA was repeated with a spatial correction added in the form of a conditioning (Z) matrix composed of Moran’s Eigenvector Maps (MEMs). To create the conditioning matrix, the R package *adespatial* (Dray *et al*. 2018) was used to construct a distance-based spatial weighting matrix from which the *mem* function was used to generate the MEMs for the 19 Atlantic Canada sampling locations. The first two MEMs were included with the genetic and environmental data on a per-individual basis and then specified as the conditioning matrix within the RDA.

Lastly, we repeated the RDA analysis on a per location basis (using minor allele frequencies for each sampling location as the genetic information) to see if the relationship of genetics and environment varied when inspected on different scale. To do this the same random 1% subsample of all MAF values used in the correlation analysis was used as the response matrix and joined to the set of uncorrelated environmental variables on a per location basis. The redundancy analysis and assessment of SNPs associated with environmental variation was repeated using the same methods as described for the first per-individual RDA.

## RESULTS

### Draft Reference Genome Assembly and quality assessment

The cunner genome assembly used in this study (NCBI accession number: GCA_020745685.1) contained 0.72 Gbp of sequence, across 24 chromosomes and 39 additional unplaced scaffolds. Other species from the Labridae family have reference genomes with 24 chromosomes, suggesting that there is genome wide coverage for cunner (Lie *et al*. 2018; Mattingsdal *et al*. 2018; Liu *et al*. 2021). Overall, the GC content of the cunner genome assembly is 0.413, and the missing nucleotide frequency was 0.0058. BUSCO analysis identified: 3576 (98.3%) complete (3544 (97.4%) single-copy, 32 (0.9%) duplicated), 20 (0.5%) fragmented, and 44 missing BUSCOs. This result further suggests a high-quality genome assembly. The mitochondrial genome was also characterized, and it contained 16,494 bp with no missing nucleotides and GC content of 0.4825.

A total of 15.41% (112.8Mb) of the 0.72 Gbp cunner genome was annotated as repetitive elements; of this, transposons accounted for 7.04% of the genome (2.76% class I TEs, and 4.28% class II TEs). The repetitive content of the cunner genome falls at the lower end of the range of values reported for sequenced Labridae genomes, with less repetitive content than corkwing wrasse (*Symphodus melops*; 18.8% of 0.61 Gbp genome), ballan wrasse (*Labrus bergylta*: 28.5% of 0.81 Gbp genome), and humphead wrasse (*Cheilinus undulatus*; 46% of 1.2 Gbp genome) (Lie *et al*. 2018; Mattingsdal *et al*. 2018; Liu *et al*. 2021).

Alignment of the cunner genome to the humphead wrasse (GCF_018320785.1) and New Zealand spotty wrasse (GCF_009762535.1) genomes using megablast and a percent identity threshold of 80% provided further evidence of a high-quality genome assembly, in both cases yielding 24 pairs of homologous chromosomes (Figure 2; Figure S1; Figure S2). This suggests that the genomic architecture of the Labridae family is conserved, with no evidence of Robertsonian fusion or fission events across the three chromosome level genome assemblies examined. The prevailing linear nature of the alignment blocks suggests strong synteny between the different genome assemblies, providing no evidence of either chromosome rearrangements or large-scale genome assembly errors.

**Figure 2.**
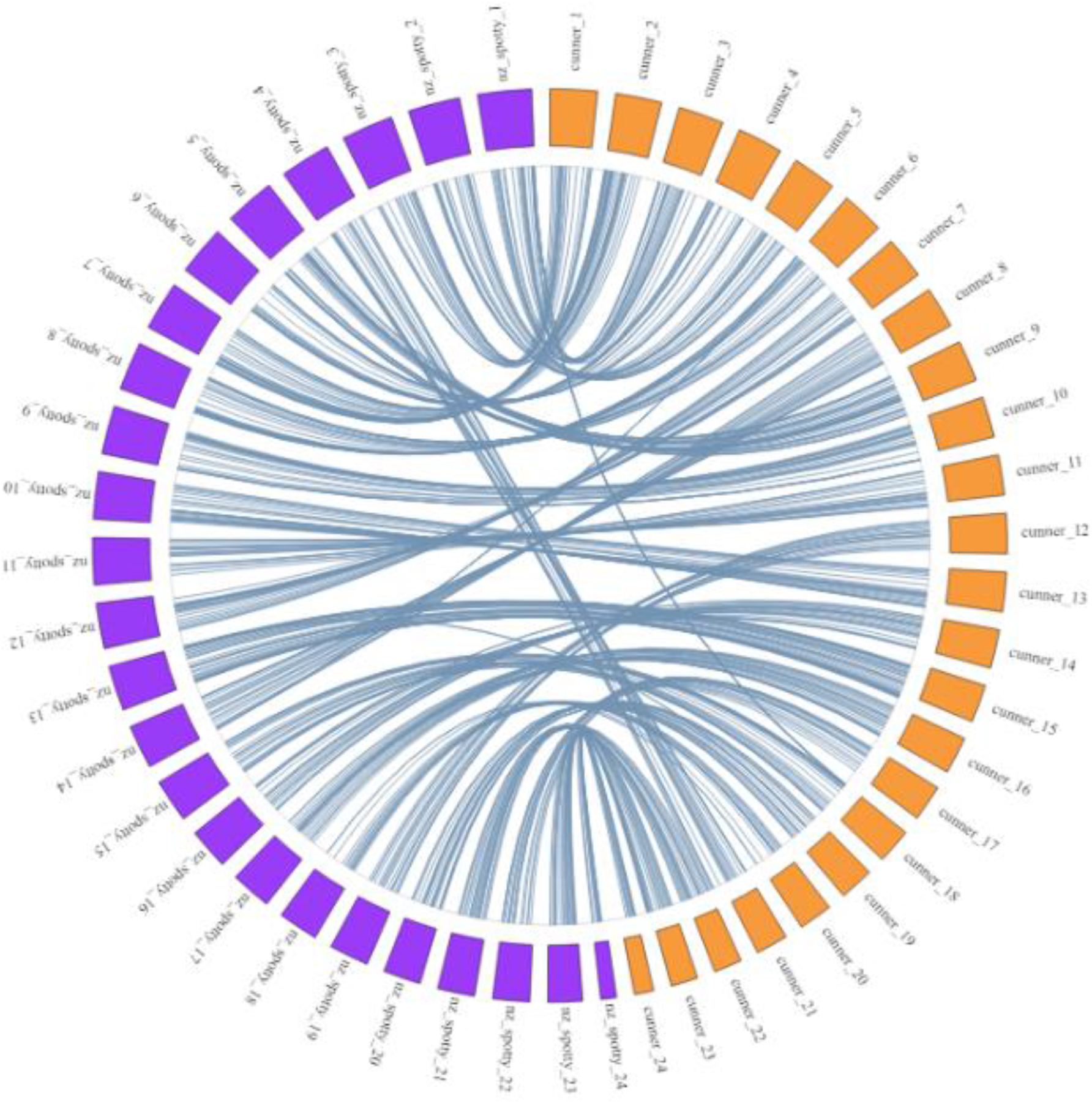
Circos plot displaying the alignment of the cunner genome to the New Zealand spotty wrasse genome. The orange rectangles represent the 24 chromosomes of cunner and the purple rectangles represent the 24 chromosomes of New Zealand spotty wrasse, both of which are numerically ordered from the top to the bottom of the plot. The blue lines indicate homologous regions of the two genomes identified via the megablast alignment of the cunner genome (query) to the New Zealand spotty wrasse genome (subject). Alternative visualization of the data can be found in Figure S2.

### Whole genome resequencing and characterizing genomic variation

The whole genome resequencing pipeline yielded 11,574,435 molecular markers for the 805 wild cunner individuals. The average depth of sequencing for the 751 Atlantic Canada individuals was 13.95x (standard deviation = 6.52), while the 54 samples from Tuckerton NJ (TUK) displayed lower depth of coverage with an average of 2.40x (standard deviation = 0.73). There are several possible reasons for the reduced coverage within the TUK samples, the most likely seems to be that the samples were smaller (collected as larvae in plankton sampling) and older (collected in 2012) so DNA degradation may have resulted in decreased sequencing yields.

The PCA of the complete set of 805 individuals showed that principal component 1 (PC1 = 5.70%) explained the largest proportion of variation and separated the TUK samples from the Atlantic Canada samples (Figure S3). Along the secondary axis of variation, there was evidence of clusters forming within the Atlantic Canada samples. The PCA of genomic variation for only the 751 Atlantic Canada samples was conducted to resolve population structure in this region. This PCA resulted in first and second principal components (PCs) that explained 0.64% of the genomic variance and 0.23% of genomic variance respectively (Figure S4). PC1 provided the most interpretable structure to the data; individuals from the same sampling locations tended to cluster together on PC1, suggesting this axis was in some manner related to geography. Along the PC2 axis, there were two outlier individuals from the Alderney (ALD) sampling location identified that were determined to be duplicate samples and, conservatively, were removed prior to the *k*-means clustering and other analyses (additional screening revealed no evidence of additional duplicate samples). Subsequent *k*-means clustering of the PCs for the 749 individuals resolved individuals into four groups that had distinct geographic distributions (Figure 3; Figure S5). Based on the *k*-means clustering of their genomic PCs, the Atlantic Canada cunner samples were assigned to four populations: Nova Scotia & New Brunswick (NSNB), Northwest Newfoundland (NWNL), Northeast Newfoundland (NENL), and Southeast Newfoundland (SENL) (Figure 3).

**Figure 3.**
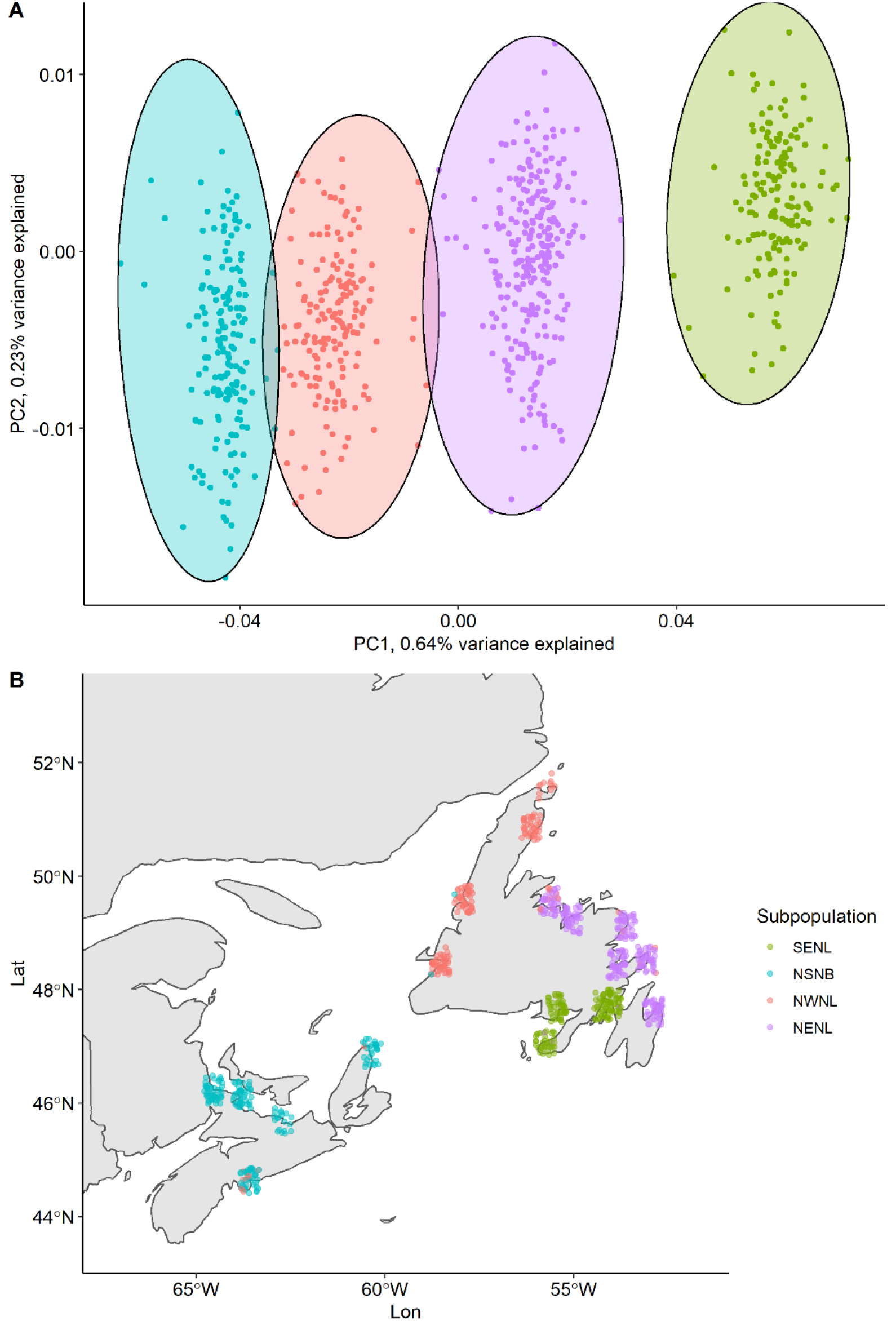
**A)** Scatter plot of Principal Components (PCs) of genetic variation for 749 cunner individuals from Atlantic Canada. The coloured ellipses denote the genetic populations identified through *k*-means clustering of PCs and the cluster to which individuals were assigned. **B)** Map of sampling locations where colour-coordinated points represent individuals and the subpopulation to which they were assigned in the *k*-means clustering.

### Characterizing genomic differentiation

Comparison of the Canadian subpopulations to the New Jersey (NJ) samples demonstrated elevated genome wide *F*_ST_ (Table 2), suggesting high genetic divergence of the NJ samples from the Atlantic Canada subpopulations. Across the pairwise comparisons of the Atlantic Canada populations, the genome-wide, per-loci *F*_ST_ values were low, with a mean of genome-wide *F*_*ST*_ = 3.2e-3 for all the pairwise comparisons (Table 2). Distinct peaks were observed within the pairwise comparisons, and the distribution of high *F*_ST_ loci deviated from expected for selectively neutral loci, with clustering of excessive numbers of outliers providing evidence of signatures of selection. The number and magnitude of *F*_ST_ peaks in the pairwise comparisons appeared to relate to the location of the different populations, although not directly to their geographic distance from one another (Figure 4; Table 2; Table S1). The highest levels of genetic differentiation were seen between the NSNB and SENL populations, with 7837 markers displaying *F*_ST_ greater than 0.15 as well as the highest genome-wide *F*_ST_ amongst the pairwise comparisons (*F*_ST_ = 0.005886). The adjacent NENL and SENL showed the lowest divergence of any comparisons with only 71 markers with *F*_ST_ greater than 0.15.

**Table 2.**
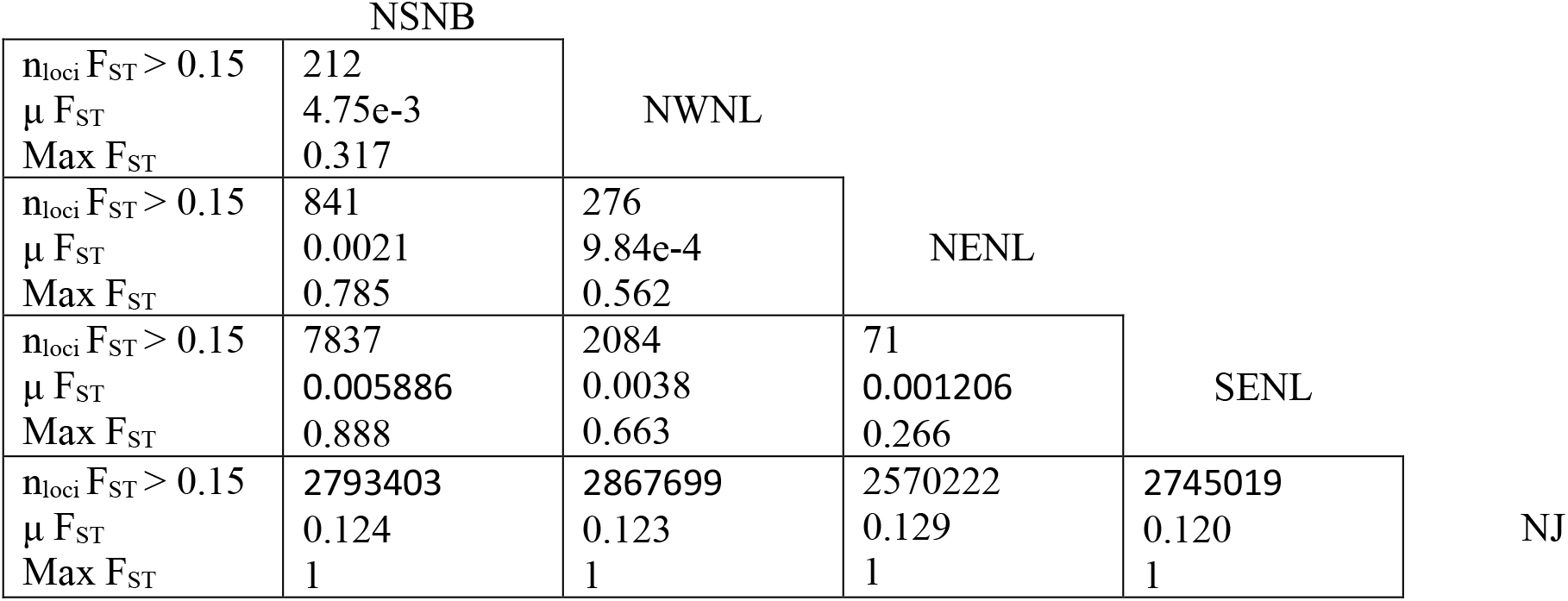
Summary statistics of the genomic differentiation (*F*_ST_) for all pairwise comparisons of cunner populations. The table gives the number of loci with an F_ST_ value greater than 0.15 (n_loci_ F_ST_ > 0.15), the mean genome wide F_ST_ for a given comparison (μ F_ST_) and the maximum F_ST_ observed for the comparison (Max F_ST_) for all of the pairwise comparisons of the four Atlantic Canada populations: Nova Scotia (NSNB), Northwest Newfoundland (NWNL), Northeast Newfoundland (NENL), and Southeast Newfoundland (SENL), as well as the set of samples from New Jersey (NJ). The vertical and horizontally aligned population designations indicate the pairwise comparison of populations that corresponds to a given cell.

|  |  |  |  |  |  |
| --- | --- | --- | --- | --- | --- |
|  | NSNB |  |  |  |  |
| $n_{loci} F_{ST} > 0.15$ | 212 | NWNL | | | |
| $\mu F_{ST}$ | 4.75e-3 | | | | |
| Max $F_{ST}$ | 0.317 | | | | |
| $n_{loci} F_{ST} > 0.15$ | 841 | 276 | NENL | | |
| $\mu F_{ST}$ | 0.0021 | 9.84e-4 | | | |
| Max $F_{ST}$ | 0.785 | 0.562 | | | |
| $n_{loci} F_{ST} > 0.15$ | 7837 | 2084 | 71 | SENL | |
| $\mu F_{ST}$ | 0.005886 | 0.0038 | 0.001206 | | |
| Max $F_{ST}$ | 0.888 | 0.663 | 0.266 | | |
| $n_{loci} F_{ST} > 0.15$ | 2793403 | 2867699 | 2570222 | 2745019 | NJ |
| $\mu F_{ST}$ | 0.124 | 0.123 | 0.129 | 0.120 | |
| Max $F_{ST}$ | 1 | 1 | 1 | 1 | |

**Figure 4.**
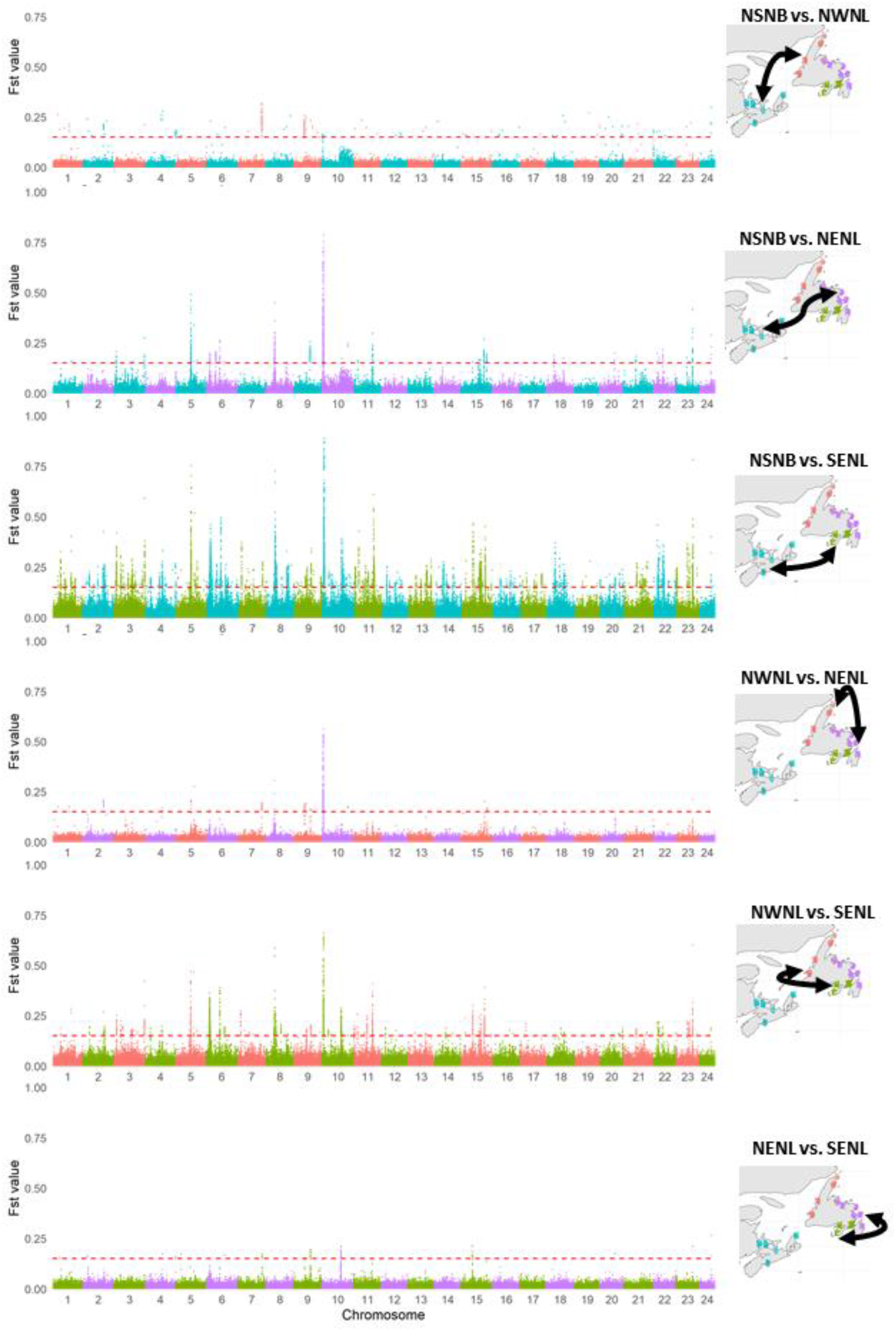
The genomic differentiation of all pairwise comparisons of cunner populations. Manhattan plots of the per locus pairwise *F*_ST_ comparisons for the Atlantic Canada subpopulations. The dashed red line indicates an *F*_ST_ threshold of 0.15. The alternating colours indicate the breakpoints between chromosomes, and the text and map to the right of a given Manhattan plot indicates the pairwise comparison shown.

A search of the genome wide *F*_ST_ data of the Atlantic Canada comparisons for significance peaks identified 241 in total across the six pairwise comparisons (Table S1; Figure 4). There were several regions of low differentiation identified: chromosome 19 possessed no *F*_ST_ peaks across any comparisons and chromosomes 13, 16, 17, and 20 contained peaks in only the NSNB to SENL comparison.

Chromosome 10 had the most prominent differentiation signal, with peaks recorded for each of the six pairwise comparisons and peaks in matching locations in four of the six pairwise comparisons. There were additional synonymous *F*_ST_ peaks observed on chromosomes 8, 9, and 15, and to a lesser extent on chromosomes 7, 11, and 23.

The reduced PCA utilizing only outlier loci (n = 7698) from within the 241 *F*_ST_ significance peaks revealed that these loci explained a larger portion of shared variation characterizing population genetic clusters than when using the whole genome data and that the same relationship of the subpopulations was identified, albeit with less distinct boundaries between the subpopulations (Figure S6). The first and second PC axes explained 14.72% and 3.06% of the variance respectively (Figure S6). The control analysis of 8000 random markers from outside of the adaptive peaks, explained less variance (PC1 = 0.649%, PC2 = 0.257%), but produced the same relationship of the subpopulations in the PC space (Figure S7).

### Scan for evidence of selection

The selection scan revealed evidence of selection in the form of regions of the genome with XP-nSL outlier values (defined as 100 Kb windows containing a locus with an XP-nSL score ≥10 or ≤-10) across the pairwise comparisons (Figure S8). Additionally, these signatures of selection were consistently observed in regions of the genome characterized as *F*_ST_ peaks. The strongest signatures of selection were observed on chromosome 8 (at 8.1Mb) and on chromosome 10 in the region of 1.2-1.5Mb (Figure 5), both of which were identified in four of the six pairwise comparisons and aligned with observed *F*_ST_ peaks (Figure 4; Figure 5; Figure S8). Other regions with evidence of selection (on chromosomes 4, 5, 9, 10, 11, 15, and 23) also co-localized with *F*_ST_ peaks (Figure 4; Figure S8).

**Figure 5.**
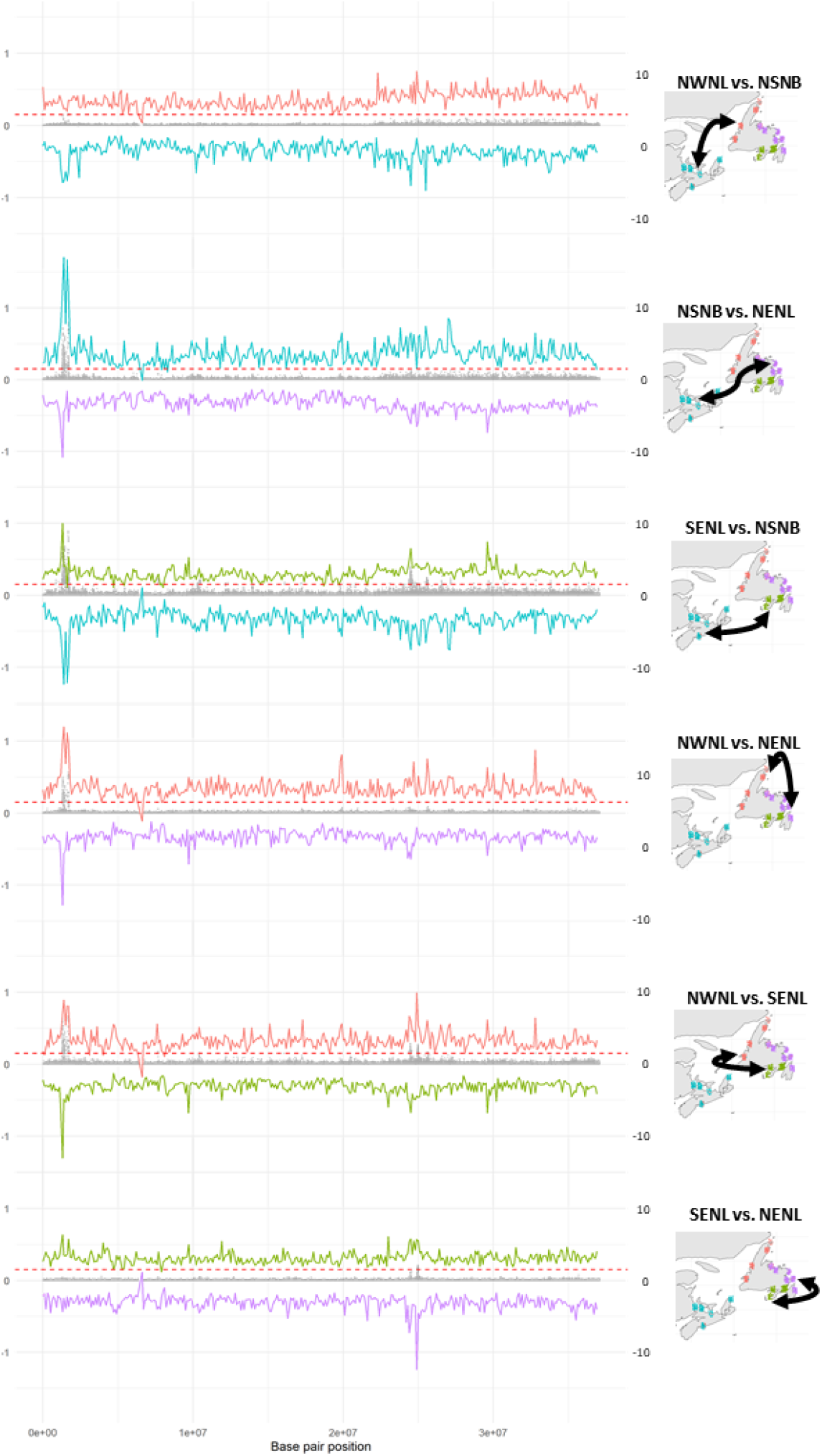
Evidence of differentiation and selection on cunner chromosome 10 for all pairwise comparisons of cunner populations. The grey dots indicate the per locus pairwise *F*_ST_ comparisons for the Atlantic Canada subpopulations. The y axis on the left-hand side of the plots relate to the *F*_ST_ values and the y axis on the right side of the plots relate to the XP-nSL scores. The dashed red line indicates an *F*_ST_ threshold of 0.15. Chromosome 10 minimum and maximum normalized XP-nSL scores for 100kb windows of each pairwise comparison of Atlantic Canada populations. The top and bottom lines indicate the minimum and maximum XP-nSL scores respectively. The minimum and maximum for each pairwise comparison indicates the population that the XP-nSL values relate to (*i.e*. in the first comparison NSNB vs. NWNL, the maximal XP-nSL is evidence of selection in NWNL and is therefore red, while the minimum XP-nSL is evidence of selection in NSNB and is therefore teal).

### Evidence of reduced recombination

Calculated LD scores were queried to search for areas of reduced recombination. These results were compared to genomic differentiation and selection results to search for evidence of population-specific structural variants associated with adaptive divergence of the populations. None of the Atlantic Canada populations displayed evidence of population-specific regions of elevated LD. Across populations, there was recurring evidence of elevated per-window median *r*^2^ (>0.05) at several regions of the genome, specifically on chromosomes 5, 13, and 18 (Figure S9). This suggests reduced recombination or structural variants in these genomic regions across the sampled range of cunner. There was no evidence of the elevated LD regions being associated with signals of adaptive divergence; only the LD peak on chromosome 5 was on the same chromosome as a recurring signal of adaptive divergence, but the location of elevated LD was greater than 4Mb from the observed *F*_ST_ significance peak.

### Association of genetic and environmental variation

The environmental data for the 19 Atlantic Canada locations featured numerous highly correlated variables. PCA allowed for these correlated variables to be summarized along their principal axes of variation. Assessment of the correlation of the principal components with the input environmental variables allowed for the environmental variables significantly contributing to the axes of variation to be identified. The first five principal component axes explained 88.95% percent of the cumulative variance for the environmental values (PC1= 48.30% variance explained, PC2 = 19.67%, PC3 = 9.81%, PC4 = 6.36%, PC5 = 4.81%). The first PC, explaining almost half of the total environmental variation, displayed significant correlations with many input variables; there were significant r^2^ values greater than 0.85 for each of the following inputs: *BO2_dissoxrange_bdmax, BO2_dissoxmean_bdmax, BO2_temprange_bdmax, BO2_tempmin_bdmax, BO2_tempmean_bdmax, BO2_tempmax_bdmax, BO2_icecovermean_ss, BO_calcite*, and less than -0.85 (a negative correlation of equal magnitude) for: *BO_dissox, BO2_salinitymean_bdmax* (Bosch *et al*. 2017), indicating that these predictors significantly contributed to this PC of environmental variation. These predictors suggests that the primary axes of environmental variation was related to temperature and oxygen range at the sea bottom.

There was a significant correlation observed between PC1 of the environmental variation and PC1 of per-individual genetic variation from within the *F*_ST_ peaks (n_loci_= 7698) for the Atlantic Canada samples (Figure 6). This correlation was best explained by linear model with a second order (quadratic) term for the environmental PC1 variable (*r*^2^=0.5579, p <2.2e-16). The correlated principal axes of environmental variation and per-individual genetic variation suggest that genetic variation is organized along an environmental gradient that, on a geographic scale, progresses from Nova Scotia around the North of Newfoundland to its southeast coast (Figure 4; Figure 6A). A quadratic correlation between PC1 of the environmental variation and PC1 of per-individual genetic variation was also observed using the complete marker set of the Atlantic Canada samples (*r*^2^ = 0.60, p < 2.2e-16), which showed that this correlation of genetic differences and environmental variation was detectable using the complete genome as well as when the analysis considered the 7698 markers from the adaptive peaks (Figure S10).

**Figure 6.**
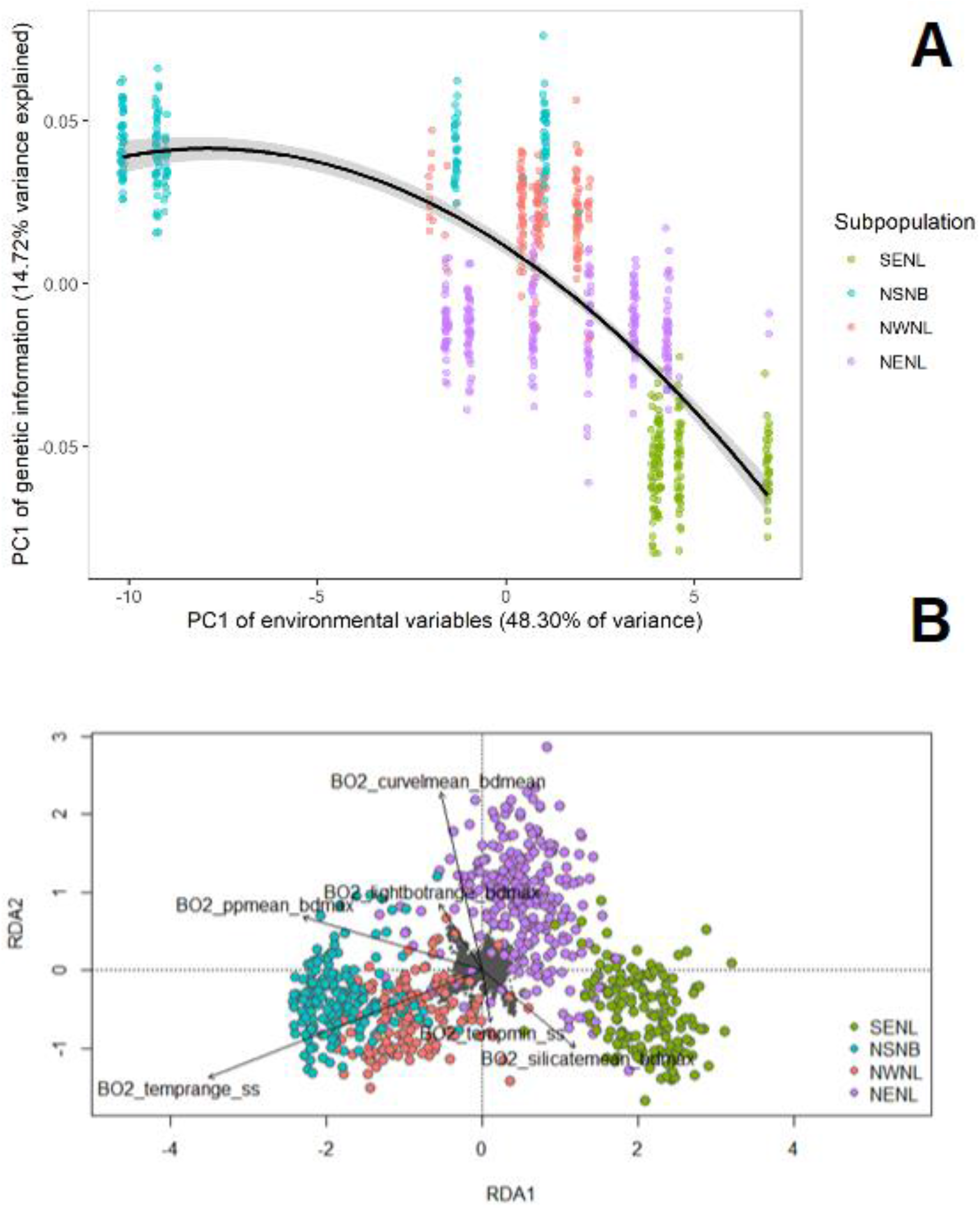
A) Scatter plot showing the relationship of the principal axes of environmental and genomic variation. Each point is an individual, and these are coloured by the subpopulation assignment for their location of origin (see Table 1). The x-axis is PC1 of environmental variation across the sampling locations (48.3% variance explained) and the y-axis is per-individual PC1 of genetic variation for the markers found within the *F*_ST_ peaks (n_loci_= 7698, 14.72% variance explained). The quadratic line of best fit is shown in black, with 95% confidence intervals in the flanking grey, the *r*^2^ of the correlation was 0.5579 (p < 2.2e-16). A version of the plot using PC1 of the complete set of genetic markers can be found in Figure S7. B) Ordination plot of the per-individual (n=749) RDA that used the six uncorrelated environmental variables (Table S3) as a predictor matrix and the genotype probabilities of the 7,698 markers from within the identified *F*_ST_ peaks as the response matrix. No correction for spatial structure was here applied (see Figure S8 for spatial correction version). The adjusted *r*^2^ of the model was 0.041, suggesting the constrained ordination explained approximately 4.1% of the genetic variation. Note that this value is likely an underestimate, as to increase resolution the response matrix was one-hot encoded (three columns per marker with the unique genotype probabilities) as opposed to dosage encoded. The small grey dots represent the genotype probabilities (response) and the larger points represent the individuals coloured by their population of origin. The black arrows represent the uncorrelated environmental predictors. The relative arrangement of the points and arrows on the plot represents their relationship with the RDA ordination axes (RDA1 = 0.7958 proportion variance explained, RDA2 = 0.083 proportion variance explained).

The PCA of per-location MAF explained 13.25% and 12.5% of variation respectively along PC1 and PC2. The primary axes of variation (PC1) appeared to exclusively separate the Raleigh Newfoundland population (part of the NWNL subpopulation) from the other locations. This population’s separation along the primary axes of variation for MAF is likely a result of a smaller sample size (n = 11) relative to the other locations (Table 1). The second PC displayed a significant linear correlation with PC1 of environmental variation (Figure S11). The strong correlation of environmental and genetic variation is therefore still detected if genetic variation is considered on a per population, as opposed to per individual, basis.

The relationship of genetics and environment for the Atlantic Canada sampling sites was then re-examined using a series of redundancy analyses. The selection of uncorrelated (r^2^ < 0.7) environmental variables yielded six values for use as the predictor matrix: *BO2_curvelmean_bdmean, BO2_lightbotrange_bdmax, BO2_ppmean_bdmax, BO2_silicatemean_bdmax, BO2_tempmin_ss, BO2_temprange_ss*. The RDA of per-individual genetic variation for markers within the *F*_ST_ peaks was first performed, and the adjusted r^2^ of the model was 0.041, suggesting that 4.1% of the genetic variation in the model was explained by the environment (Figure 6B) The model displayed a pattern similar to the one observed in the correlation of principal components, with individuals associating along the same environmental gradient (Figure 3; Figure 6). The first two RDA axes explained 79.6% and 8.3% of variation, with subsequent axes showing reduced importance. Inspection of RDA loading values showed only a single marker on chromosome 10 with a loading for RDA1 more than 3 standard deviations from the mean, lowering the stringency of this test to 2.5 standard deviations showed 34 markers potentially associated with this RDA axis, 31 of which were located on chromosome 10, one from chromosome 7 and one from chromosome 5.

Repeating the per-individual analysis with MEM-based spatial correction led to a reduction in the adjusted *r*^2^ of the model to 0.01; the magnitude of the separation of populations also decreased, but the pattern of the clustering of individuals from the same population were still observed in the ordination plot when spatial structure was accounted for (Figure S12). The first two axes of variation explained a smaller proportion of variation (RDA1 = 56.47%, RDA2 = 14.04%). Relative to the non-spatially corrected plot, there was a change to the markers displaying association with the primary RDA axis. When the spatial correction was applied, 239 makers had RDA1 loadings greater than 3 standard deviations from the mean, and notably this included 94 loci from chromosome 7 and 93 loci from chromosome 9 (with no loci from chromosome 10). Overall, these analyses further support the adaptive divergence of cunner along an environmental gradient and give some further indication of the relationship of the regions of selection relative to environmental conditions.

## DISCUSSION

Adaptation to ocean climate is increasingly recognized as an important driver of diversity in marine species (De Faveri *et al*. 2013; Duranton *et al*. 2018; Jahnke *et al*. 2018; Stanley *et al*. 2018; Knutsen *et al*. 2022; Pratt *et al*. 2022) and understanding of the evolutionary and ecological process involved is increasingly central to management and conservation action (Lehnert *et al*. 2019; Pratt *et al*. 2022). Here we developed the first genomic resources for the temperate reef fish, cunner, by producing a chromosome-level genome assembly, demonstrating the high quality of the assembly through comparison to closely relates species, and identifying millions of genetic variants for cunner. This information was then used to characterize the genetic structure of wild populations throughout Atlantic Canada and New Jersey, identify regions of genomic differentiation among populations, and associate genomic variation with an environmental gradient. Our analysis identified four distinct regional populations throughout Atlantic Canada consistent with fine scale geographic structuring despite pelagic egg and larval stages in this species. The genomic distribution of differentiation, genomic signatures of selection, and environmental association analysis all suggest adaption to regional differences in ocean climate is a strong driver of differentiation across the study area. This study extends a growing body of genomic analyses supporting a role for ocean climate in driving fine geographic scale differentiation in marine taxa despite apparent high dispersal potential during the early life history (Lehnert *et al*. 2018; Stanley *et al*. 2018; Watson *et al*. 2021; Knutsen *et al*. 2022). These results suggest that climate associated adaptation has driven the evolution of regional diversity in cunner, despite the potential for wide dispersal and reveal how the interplay of climate, adaptive diversity, and life history can structure marine populations.

### Population structure and cunner life history

Our clustering analyses revealed four genetically distinct cunner populations that were regionally distinct and strongly suggestive of fine scale population structure throughout Atlantic Canada. The strong signal of population structure is surprising given the early life history of cunner, but the findings closely resemble those of other Labridae. Geographically and genetically distinct populations have previously been identified in Goldsinny wrasse *(Ctenolabrus rupestris*), a closely related Labridae species found along the Atlantic coastlines of Europe and North Africa, as well as the Sea of Marmarra, the Black Sea and parts of the Mediterranean Sea (Jansson *et al*. 2020). Furthermore, work on corkwing wrasse (*Symphodus melops*) has identified a major genetic break between Scandinavia and more southern populations (Knutsen *et al*. 2013) despite a seemingly continuous marine environment. This population structure was attributed to the corkwing wrasse’s requirement for rock substrate and its short pelagic larval phase of 2-3 weeks (Knutsen *et al*. 2013). These concordant trends of regional population structure for multiple related species found on both sides of the Atlantic Ocean suggest that geographic structuring may not be species-specific and instead be common among Labridae species, possibly result from common life history patterns.

The four genetically distinct cunner populations we have identified in Atlantic Canada may be the result of adaptation to local habitat differences that result in increased survival rates and recruitment in the presence of dispersal and gene flow. As adults, cunner are found primarily in nearshore reefs, but the species has a pelagic egg and larval stage (lasting three weeks) that facilitate wide dispersal (Levin 1996; Juanes 2007). Larval dispersal is followed by a three-to-four-week settlement period, in which cunner use the structure of their habitat for shelter against predation and as refuge for their nightly period of torpor (Juanes 2007). Research on Nova Scotia cunner has shown that the survival of the settlement period correlates with habitat complexity (Tupper & Boutilier 1997). Cunner larvae may disperse widely, but their adaptive genetic predisposition to success in a habitat may strongly impact their recruitment (*i.e*. regional differences in the type of shelter available and also physiological or behavioural differences that are conducive to survival in a given type of shelter). This is supported by examination of the subpopulation assignments of individual samples relative to their sampling locations and some evidence of apparent first-generation migrants between clusters along the Newfoundland northeast coast. Genetic cluster assignment for individuals collected at a location show strong evidence of geographic structure (96.5% average intra-location concordance), but there were also locations where individuals clustering to different genetic subpopulations are identified in sympatry (Table S2). This observation of both strong population structure driven by specific adaptive peaks in the genome and putative evidence of genetic admixture across populations suggests that dispersal may be occurring, but that migrants may face strong local selection pressure resulting in fitness reductions.

### Relation of genomic and environmental variation

The correlation of the genetic and climatic gradients supports the hypothesis that the genetic divergence of populations is associated with an environmental gradient, and likely adaptive in nature. The gradient of environmental variation appears to be a more important indicator of genetic similarity and connectivity than the geographic proximity of populations. For example, while the NSNB and SENL subpopulations are not the most geographically distant of the pairwise population comparisons, they are the most dissimilar genetically, as evidenced by both the genomic variation PCs and their pairwise *F*_ST_ comparisons (Figure 4; Table 2). They occur on the extremes of the environmental gradient, suggesting that the environmental characteristics, and not geographic distances, are the main correlate with genomic diversity (Figure 6). The main contributing variables to the primary axis of environmental variation were related to temperature and oxygen range at the sea bottom. Cunner inhabit the sea floor and enter temperature-dependent daily and seasonal states of torpor in which they bury themselves into substrate or hide in crevices (Bradbury *et al*. 1995; Moran *et al*. 2019). The primary axis of environmental variation is therefore suggestive of a relationship to cunner life history; future analysis of the genes under selection may further elucidate the mechanisms under selection and the relationship between environment, behavior, and survival.

The association of environmental and genetic gradients was repeatable across different scales (per-individual and per-location) and methods (linear modelling and RDA), suggesting the presence of a strong underlying biological signal. The additional RDA with the application of a MEM-based spatial correction reduced the strength of the association between environment and genetics. This spatial correction is likely to be overly stringent because it is reliant on a Cartesian, distance-based spatial weighting matrix (Bauman *et al*. 2018). The true connectivity of the populations is likely more nuanced than the connections in the distance-based spatial weighting matrix would indicate (Bauman *et al*. 2018). For example, the connectivity of the SENL and NENL populations is overestimated by Cartesian distances as the Isthmus of Avalon presents a barrier to migration that would greatly increase the migration distances between geographically proximal populations. The differences between the spatial corrected and non-spatial corrected RDA are nonetheless of interest. The presence of the strong environmental association with the markers on chromosome 10 when no correction is applied, and the absence of this association in the spatial corrected RDA suggests that this peak of adaptive divergence is related to cunner population structure and connectivity, possibly representing an adaptive locus favourable to survival in certain environments that has spread to adjacent populations though migration and dispersal. The results indicate the peak on chromosome 10 is associated with the environmental gradient but may also be influenced by the cunner population’s geographic structure and connectivity, while the regions of differentiation on chromosomes 7 and 9 seem to be associated with the gradient of environmental differentiation even when the population structure is accounted for.

The sampling locations employed in this study are not an exhaustive representation of the range of cunner (Moran *et al*. 2019) and key genetic differences may be found in individuals from more southern locations. Within Atlantic Canada, multiple marine species have exhibited a biogeographic break at 44.61°N (±0.25) that relates to a climatic gradient driven by seasonal temperature minima in the northwest Atlantic (Stanley *et al*. 2018). The southernmost sampling location in the analysis of Atlantic Canada population structure was Dartmouth, NS (ALD), found at a latitude of 44.663°N. The sampled range of Atlantic Canada cunner is therefore exclusively north of this multi-species biogeographic break. Our single sampling location from south of this location, Tuckerton NJ, was an extreme geographic outlier that was more than 5 degrees of latitude further south than the next closest sample and had a significantly smaller sample size. Comparisons of the four Atlantic Canadian subpopulations to the New Jersey (NJ) samples all demonstrated mean genome wide *F*_ST_ in excess of 0.12, which did not allow for fine scale differences between these southern samples and their northern counterparts to be resolved. The magnitude and genome wide distribution of Atlantic Canada differentiation with NJ suggests significant variation exists across the complete range of cunner that remains to be fully explored.

The observed genetic structuring within Atlantic Canada represents the northern extreme of the range of cunner, where severe environmental conditions may be contributing to the selective pressures and adaptive diversity. For example, cunner experience months-long physiological torpor in response to prolonged periods of low temperature (Bradbury *et al*. 1995; Moran *et al*. 2019). This is a radically different environment than the southern limits of the cunner range; the complete absence of sea ice in Tukerton NJ and habitats of similar latitudes would possibly allow cunner to entirely forgo yearly torpor cycles and the associated selective pressures (Bosch *et al*. 2017). There remains the potential for significant adaptive differentiation outside the area of sampling in the present study due to large environmental and potential life history differences. Additional sampling from an increased area could explore if the observed genetic structuring is exclusive to the northern edge of cunner’s habitable range where environmental gradients are extreme and if their dispersal-enabling life history results in more *panmictic* populations within more moderate environments.

### Relationship of adaptive divergence and observed population structure

The pairwise comparison of the populations revealed signals of adaptive divergence against a genomic background of low inter-population differentiation, which provided evidence of environmentally associated population structure. Both the number of *F*_ST_ peaks and the magnitude of co-localized *F*_ST_ peaks related to the identified fine scale population structure and environmental gradient within Atlantic Canada. There was lack of association between regions of adaptive divergence and signals of reduced recombination (LD), meaning there was no evidence to suggest that the regions under selection were associated with structural variants (SVs) (*e.g*. inversions) housing conserved polygenic haplotypes protected from recombination. The well described relationship between structural genomic variants and evolutionary adaptation (Mérot *et al*. 2020; Oomen *et al*. 2020) makes this result surprising. There are several non-biological causes that may have led to a lack of elevated LD near the adaptive peaks. If there was more pronounced LD at other neutral locations in the genome, then SVs in the regions of the adaptive peaks may have been overlooked due to not being outlier values. Secondly but perhaps less likely, these adaptive peaks may occur in regions of the genome where meiotic recombination occurs normally and may be indicative of monogenic or otherwise acute genetic differences (Mérot *et al*. 2020; Oomen *et al*. 2020).

The reduced PCA of only loci from adaptive divergence peaks showed that there was a large contribution of these loci to the genomic structure of populations, with PC1 explaining a 14.72% of variation. This was higher than the variance explained in the initial genome wide PCA, where PCs 1 and 2 explained 0.64% and 0.23% of the observed variation respectively. This suggests that most of the genome shows minimal genetic differentiation and that the adaptive peaks provide a large contribution to population differences. The relationship of the populations in the PC space remained the same as in the reduced PCA and the PCA of complete genetic variation, but the boundaries between the four populations were not as clearly resolved in the former. Work in Atlantic salmon within Atlantic Canada has also shown a pattern of low genome wide differentiation with regions of elevated differentiation (Moore *et al*. 2014). A more extreme version of this pattern has been observed in research on another marine species, Atlantic herring (*Clupea harengus*), in which a PCA utilizing 794 significant *F*_ST_ outliers explained 43% of variation along the first principal component axes (Han *et al*. 2020). Within herring, a higher amount of variation was explained by lower number of loci relative to the present study of cunner. Therefore, although the importance of the peaks of adaptive divergence in resolving cunner population structure is high, these loci do not appear to be the exclusive source of genetic differences between the populations and there are likely also neutral genetic differences that have evolved outside of the main regions of differentiation. The existence of neutral differences is supported by the control analysis of 8000 random markers from outside of the adaptive peaks, which explained less variance (PC1 = 0.649%, PC2 = 0.257%) but still revealed the same relationship spatial structure of the populations within the PC space (Figure S7). Overall, this supports the existence of strong genetic structure that is primarily, but not exclusively, the result of adaptive divergence to environmental differences.

### Conclusion

Characterization of genome wide genetic diversity of cunner samples from throughout Atlantic Canada has shown evidence of fine scale population structure and an association of genomic diversity with an environmental gradient at the northern limits of the species range. Evidence suggests that the differences between populations are adaptive responses to environmental variation, with low genome-wide divergence of populations and signals of differentiation and selection localizing to specific regions of the genome. The functions of the regions of the genome undergoing adaptive selection are not characterized at present, but given knowledge of cunner life history, an association with post-settlement survival and recruitment seems likely. As genomic resources for the species are expanded, future work examining the proximal mechanisms under adaptive selection can be explored in more detail. Expansion of the sampled range of cunner can show if the observed pattern of adaptive responses to environmental variation extends into the more southern range of cunner. The cunner population structure that we have identified within Atlantic Canada warrants consideration in aquaculture policy and management decisions that aim to promote the responsible and environmentally informed future use of cunner as a cleaner fish within Atlantic Canadian aquaculture.

## ACKNOWLEDGEMENTS

We thank Olivier Fedrigo for coordination of DNA extraction and sequencing for the genome assembly at VGP Rockefeller. Funding and support for this project was provided by Fisheries and Oceans Canada funding programs, the Aquaculture Collaborative Research and Development Program (ACRDP) in partnership with: Newfoundland Aquaculture Industry Association (NAIA), Grieg NL Seafarms Ltd., Cooke Aquaculture Inc., and Mowi Canada East, and the Program for Aquaculture Regulatory Research. Nell den Heyer assisted with collection of cunner samples in Cape Breton. Tony Einfeldt assisted with collection of cunner samples in Southern Nova Scotia. Members of the Bradbury and Bentzen labs assisted with sample collection in Newfoundland and Nova Scotia.

## AUTHOR CONTRIBUTIONS

The study was designed by IRB, TK, and CMN. Sample collection was done by TK, SJD, SJL, and BFW. Analysis design was done by CMN, TK, MKB, BLL, and IRB. Bioinformatics analyses and data visualizations were conducted by CMN. Initial manuscript preparation was done by CMN. All authors contributed to the revisions of the manuscript.

## SUPPORTING INFORMATION

Additional supporting information may be found in the online version of the article at the publisher’s website.

