## Supplementary material for "Whole genome sequencing reveals fine-scale climate associated adaptive divergence near the range limits of a temperate reef fish": S1-tables_and_figures

**SUPPLEMENTARY TABLES AND FIGURES**

**Table S1.** The number of *F*_ST_ peaks summarized by chromosome and pairwise comparison. A peak is here defined as five or more markers with greater than 0.15 *F*_ST_ and with no more than 100Kb between adjacent group members.

| Chromosome | NSNB_vs. NENL | NWNL vs. NENL | NWNL vs. NSNB | NWNL vs. SENL | SENL vs. NENL | SENL vs. NSNB |
| --- | --- | --- | --- | --- | --- | --- |
| 1 | 0 | 0 | 0 | 1 | 0 | 4 |
| 2 | 0 | 1 | 1 | 3 | 0 | 6 |
| 3 | 2 | 0 | 0 | 6 | 0 | 13 |
| 4 | 0 | 0 | 2 | 1 | 0 | 6 |
| 5 | 1 | 1 | 0 | 2 | 0 | 4 |
| 6 | 3 | 0 | 0 | 5 | 0 | 14 |
| 7 | 0 | 1 | 1 | 2 | 0 | 4 |
| 8 | 1 | 0 | 0 | 4 | 0 | 10 |
| 9 | 1 | 1 | 1 | 2 | 1 | 7 |
| 10 | 1 | 1 | 0 | 4 | 1 | 11 |
| 11 | 2 | 0 | 0 | 6 | 0 | 10 |
| 12 | 0 | 0 | 0 | 1 | 0 | 5 |
| 13 | 0 | 0 | 0 | 0 | 0 | 6 |
| 14 | 0 | 0 | 0 | 1 | 0 | 3 |
| 15 | 2 | 1 | 0 | 4 | 1 | 11 |
| 16 | 0 | 0 | 0 | 0 | 0 | 5 |
| 17 | 0 | 0 | 0 | 0 | 0 | 4 |
| 18 | 1 | 0 | 0 | 1 | 0 | 8 |
| 20 | 0 | 0 | 0 | 0 | 0 | 5 |
| 21 | 1 | 0 | 0 | 1 | 0 | 7 |
| 22 | 1 | 0 | 0 | 3 | 0 | 10 |
| 23 | 1 | 0 | 0 | 3 | 0 | 5 |
| 24 | 1 | 0 | 0 | 0 | 0 | 1 |

**Table S2.** The genomic *k*-means cluster assignments of individuals from each location. See Table 1. for the full location names corresponding to each location ID. The table is sorted by primary cluster assignment, and within each cluster longitudinally from west to east.

| Location ID | NSNB | NWNL | NENL | SENL | Latitude | Longitude | Primary subpopulation assignment |
| --- | --- | --- | --- | --- | --- | --- | --- |
| PDC | 47 | 0 | 0 | 0 | 46.2401 | -64.5286 | NSNB |
| CTR | 44 | 0 | 0 | 0 | 46.134 | -63.777 |  |
| ALD | 36 | 7 | 0 | 0 | 44.663 | -63.57 |  |
| PIC | 19 | 0 | 0 | 0 | 45.67441 | -62.7047 |  |
| WHI | 29 | 1 | 0 | 0 | 46.88637 | -60.3487 |  |
| STE | 1 | 45 | 0 | 0 | 48.51383 | -58.5378 | NWNL |
| RKH | 1 | 45 | 0 | 0 | 49.59103 | -57.919 |  |
| ROD | 0 | 40 | 0 | 0 | 50.86504 | -56.1298 |  |
| RAL | 0 | 11 | 0 | 0 | 51.56469 | -55.732 |  |
| BTN | 0 | 10 | 36 | 0 | 49.54746 | -55.6374 | NENL |
| LWP | 0 | 0 | 43 | 0 | 49.24239 | -55.0564 |  |
| BVA | 0 | 0 | 46 | 0 | 48.445 | -53.845 |  |
| VAL | 0 | 3 | 40 | 0 | 49.12228 | -53.6098 |  |
| CTB | 0 | 3 | 41 | 0 | 48.516 | -53.076 |  |
| CBS | 0 | 0 | 44 | 0 | 47.593 | -52.887 |  |
| GBA | 0 | 0 | 2 | 34 | 47.0993 | -55.7511 | SENL |
| PLC | 0 | 0 | 0 | 43 | 47.68018 | -55.4304 |  |
| BHN | 0 | 0 | 0 | 32 | 47.70929 | -54.2151 |  |
| ARN | 0 | 0 | 1 | 45 | 47.75792 | -54.00729 |  |

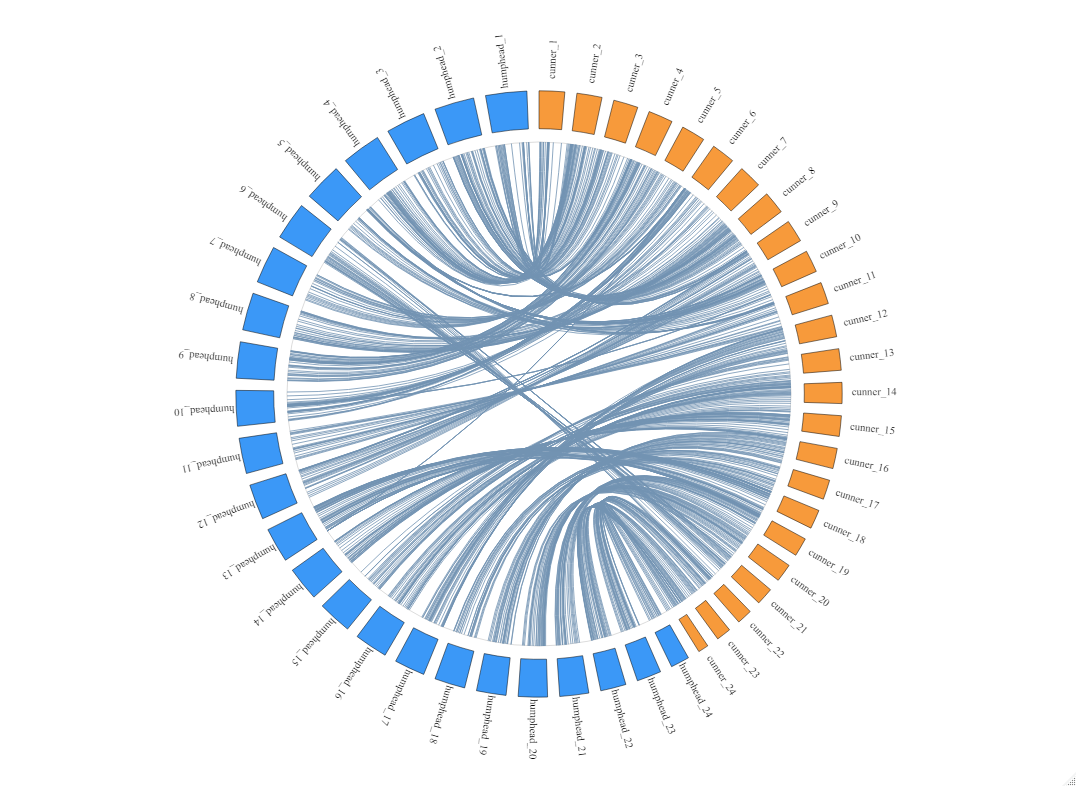

**Figure S1.** Circos plot displaying the alignment of the cunner genome to the humphead wrasse genome. The orange rectangles represent the 24 chromosomes of cunner and the blue rectangles represent the 24 chromosomes of New Zealand spotty wrasse, both of which are numerically ordered from the top to the bottom of the plot. The blue lines indicate homologous regions of the two genomes identified via the megablast alignment of the cunner genome (query) to the humphead wrasse genome (subject). Alternative visualization of the data can be found in Figure S2.

A

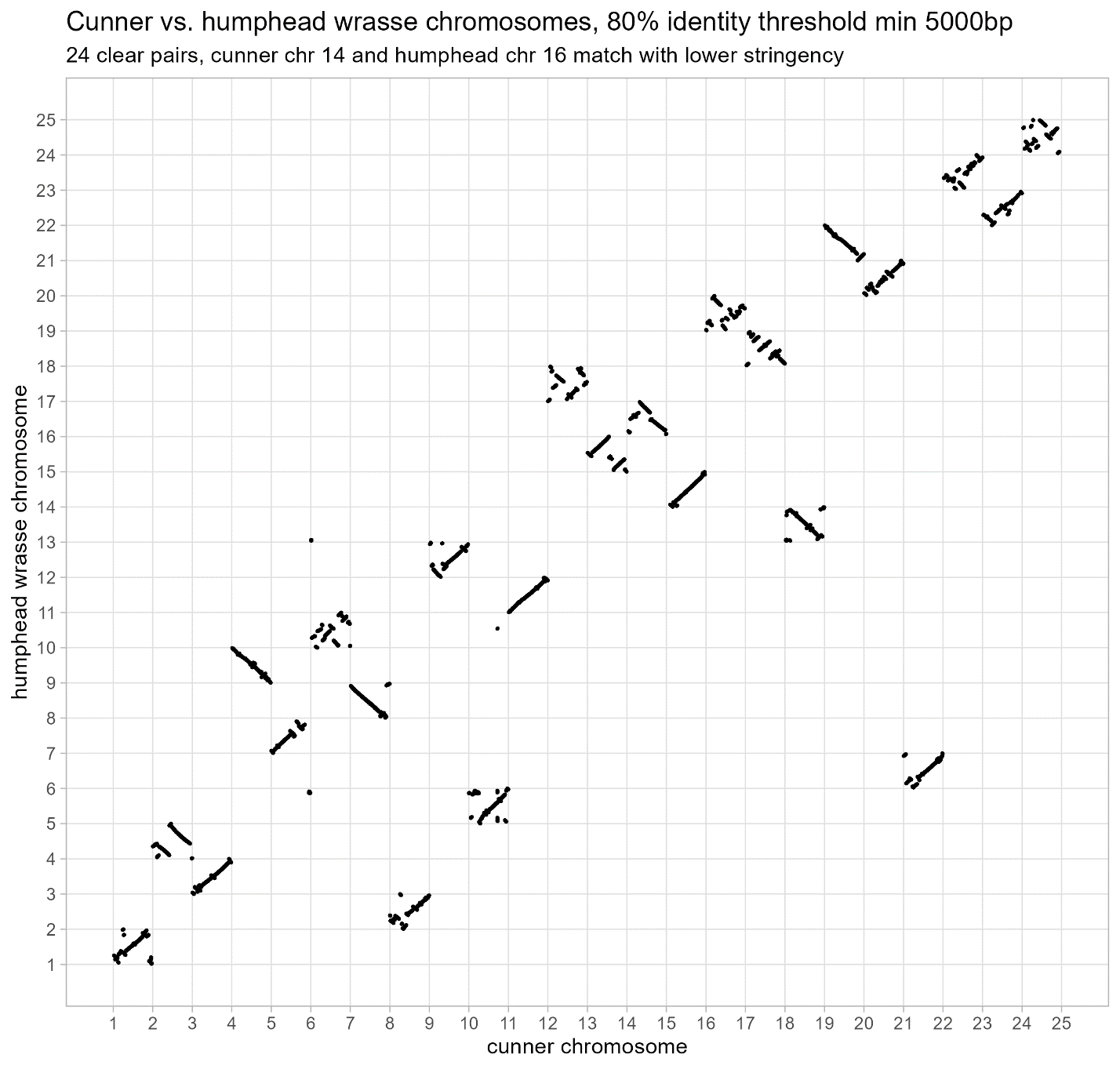

**B**

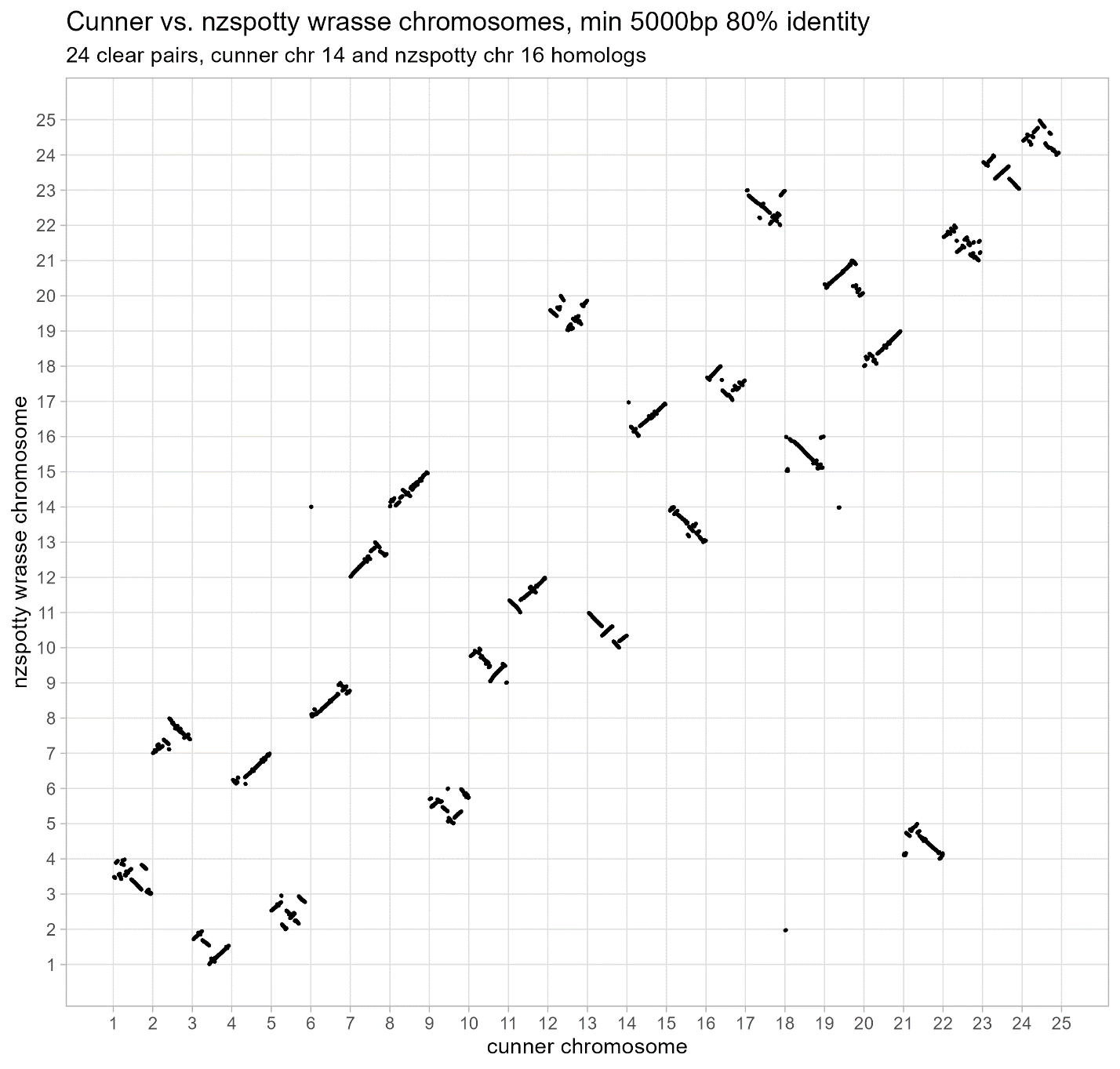

**C**

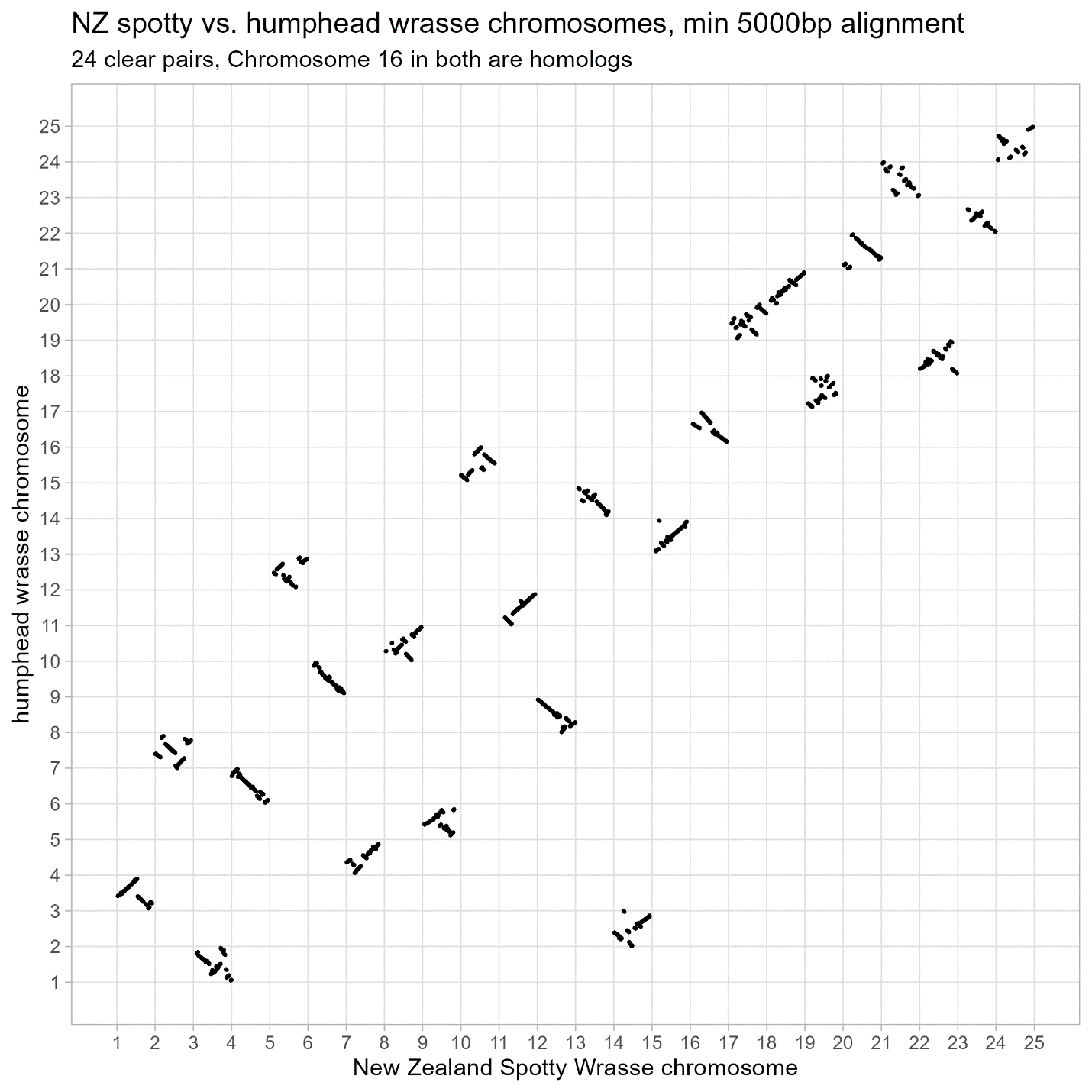

**Figure S2.** A) Dot plot of filtered megablast results for the alignment of the chromosomes of the cunner genome (query) to the chromosomes of the humphead wrasse genome (subject). B) Dot plot of filtered megablast results for the alignment of the chromosomes of the cunner genome (query) to the chromosomes of the New Zealand spotty wrasse genome (subject). **C**) Dot plot of filtered megablast results for the alignment of the chromosomes of the New Zealand spotty wrasse genome (query) to the chromosomes of the humphead wrasse genome (subject). For all plots, dots indicate megablast alignment hits (% identity >80) of greater than 5000 base pairs in length, and greater in length than 1% of both the query and subject chromosomes. The x and y label numbers intersecting the bottom left corner of a cell indicate the simplified chromosome designation of the species. Continuous lines of dots indicate strong homology between the two species chromosomes. Non-continuous sets of dots within a cell indicate either small scale chromosome rearrangements (i.e. inversions) or localized assembly errors in one or both of the genome assemblies (i.e. a contig added into the chromosome the wrong way round).

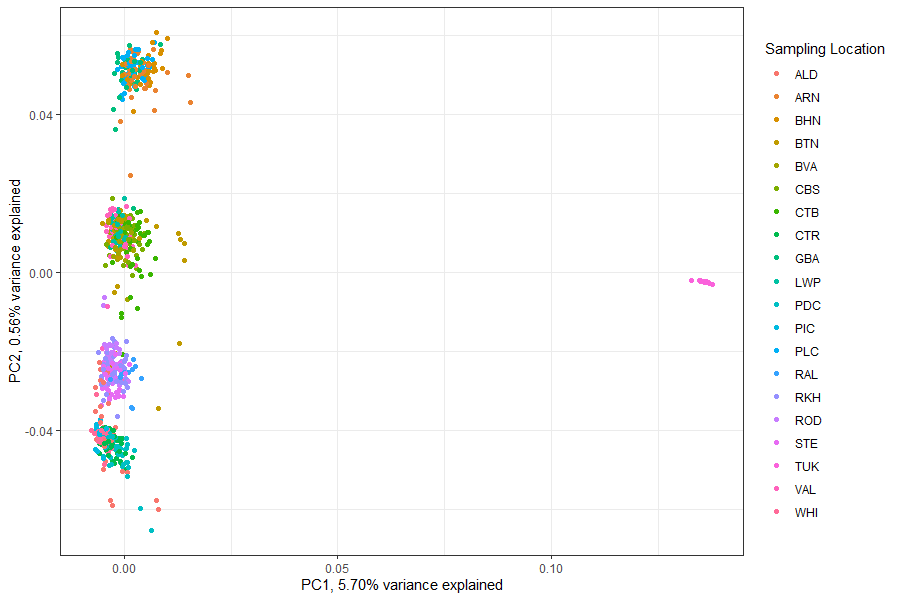

**Figure S3.** Scatter plot of the first and second PCs of the PCA of genome wide variation (11,574,435 markers) for the complete set of cunner individuals. The colours of the points indicate the population of origin for the given individuals (See Table 1 for details on the location codes). The pink points on the extreme right of the graph are the 54 samples from Tukerton, NJ (TUK).

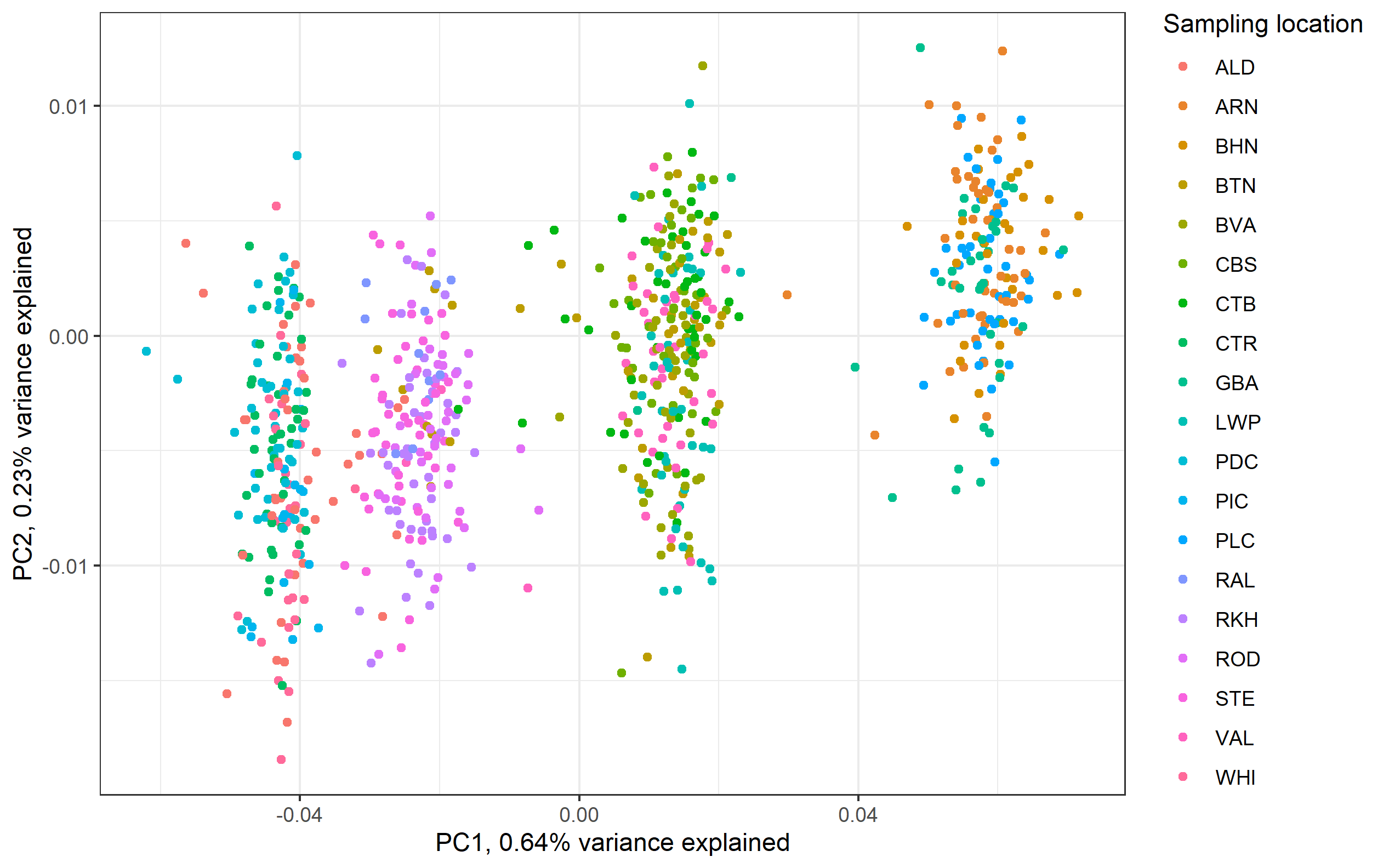

**Figure S4.** Scatter plot of the first and second PCs of the PCA of genome wide variation (11,574,435 markers) for the Atlantic Canada cunner samples (n = 749). The colours of the points indicate the population of origin for the given individuals (See Table 1 for details on the location codes).

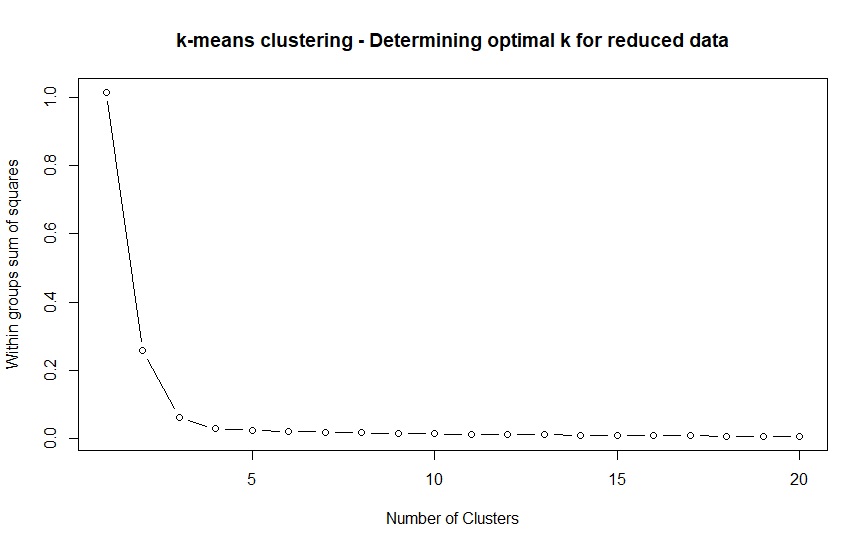

**Figure S5.** Scree plot showing the *k*-means clustering within group sum of squares for different values of *k* used to cluster the first two PCs of the Atlantic Canada samples (n=749).

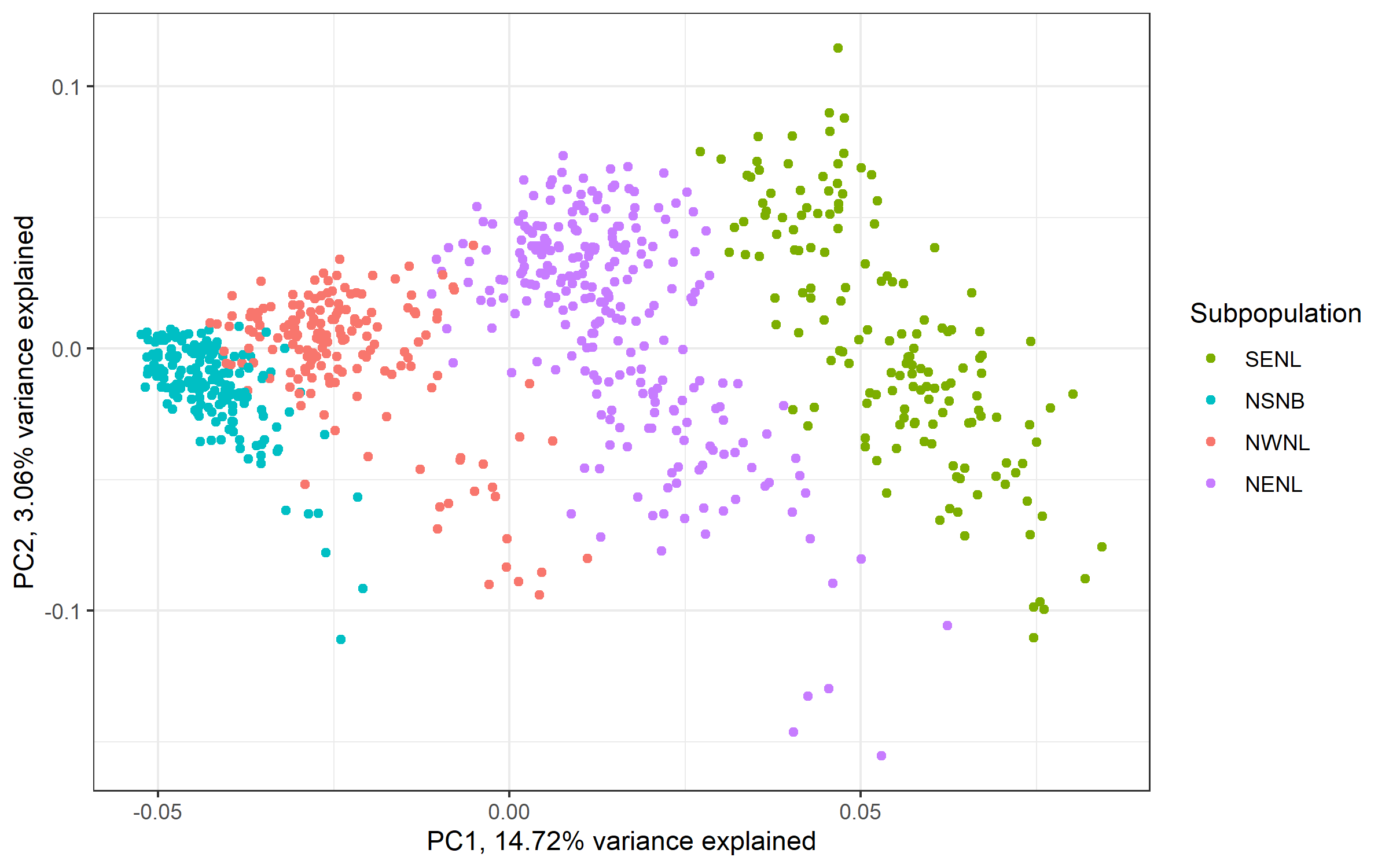

**Figure S6.** Cunner PCA of only SNPs in the *F*_ST_ peaks. Plot is per individual values for PC1 and PC2, coloured by assigned subpopulation (n_individuals_ = 749, n_loci_= 7698).

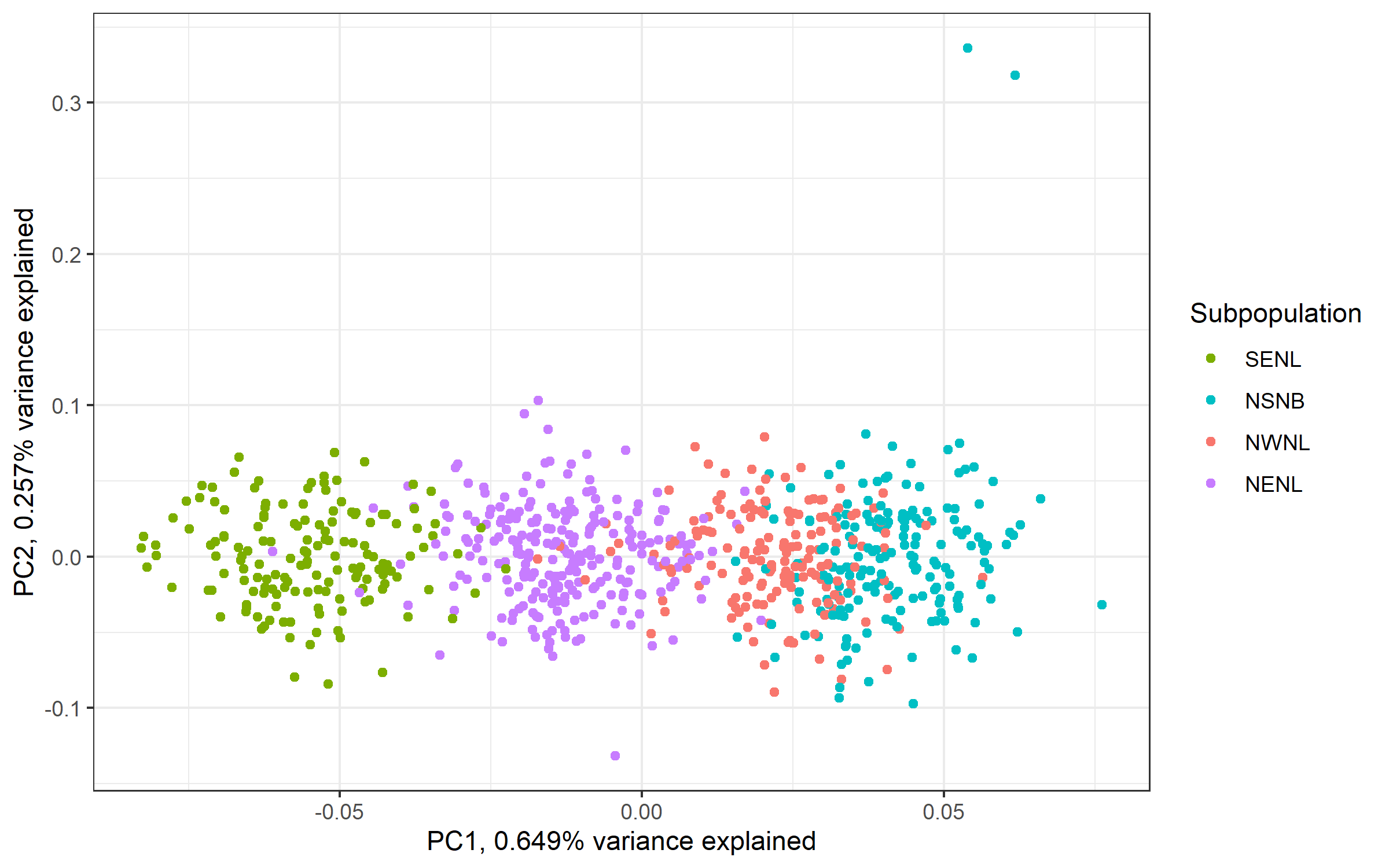

**Figure S7.** Cunner PCA of a set of 8,000 random markers selected from thoughout the genome, excluding markers found within the *F*_ST_ peaks. Plot is per individual values for PC1 and PC2, coloured by assigned subpopulation (n_individuals_ = 749, n_loci_= 8000).

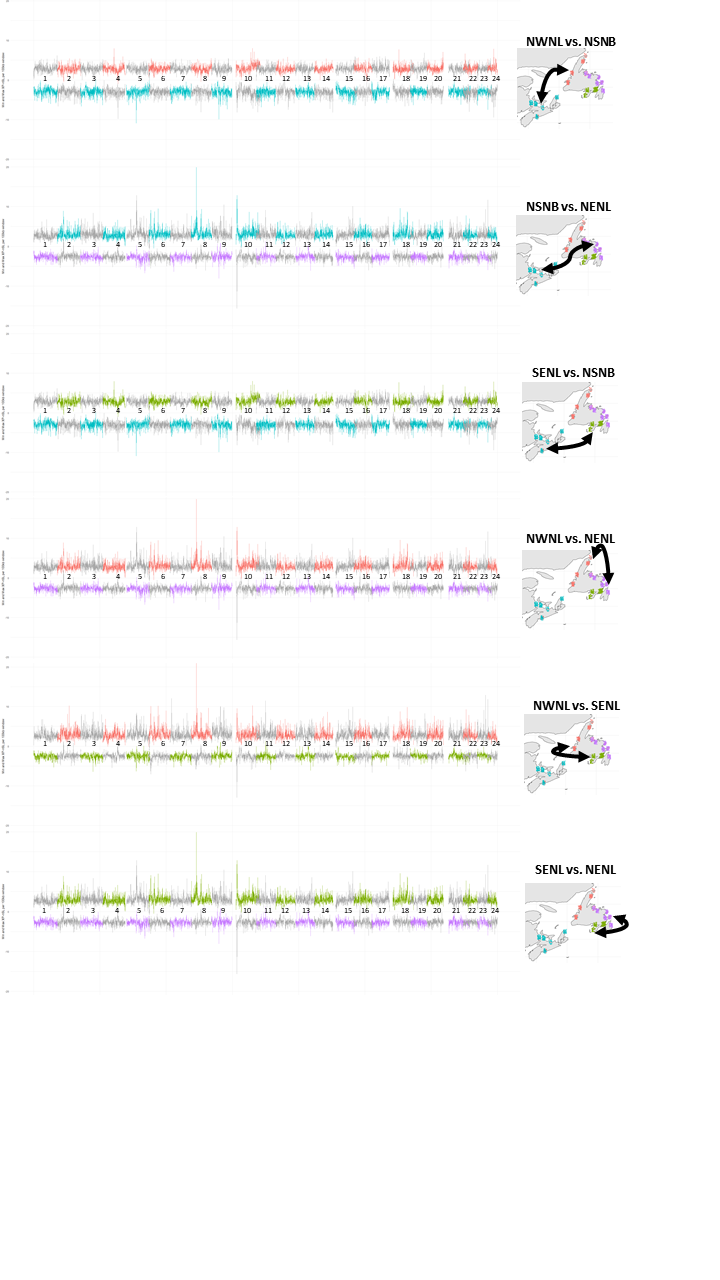

**Figure S8.** Genome wide minimum and maximum normalized XP-nSL scores for 100kb windows of each pairwise comparison of Atlantic Canada populations. The top and bottom lines indicate the minimum and maximum XP-nSL scores respectively. The alternation of grey and the subpopulation’s colour code indicate chromosome breaks and the numbers in the center of the plot indicate the chromosome number. The minimum and maximum for each pairwise comparison indicates the subpopulation that the XP-nSL values relate to (*i.e.* in the first comparison NSNB vs. NWNL, the maximal XP-nSL is evidence of selection in NWNL and is therefore red, while the minimum XP-nSL is evidence of selection in NSNB and is therefore teal).

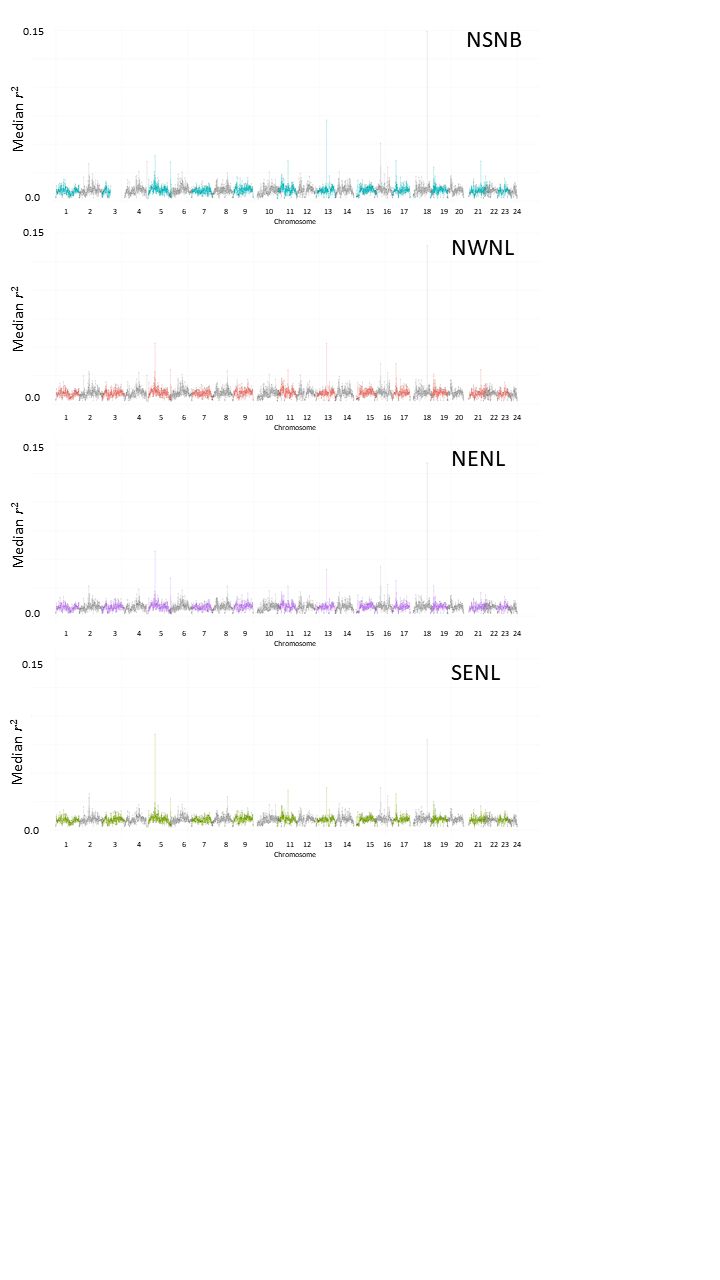

**Figure S9.** Per population Linkage Disequilibrium (LD). The four plots show the median *r*^2^ for all the comparisons involving loci from a given 100Kb window. Each locus was compared to all markers within 100Kb of its specific location.

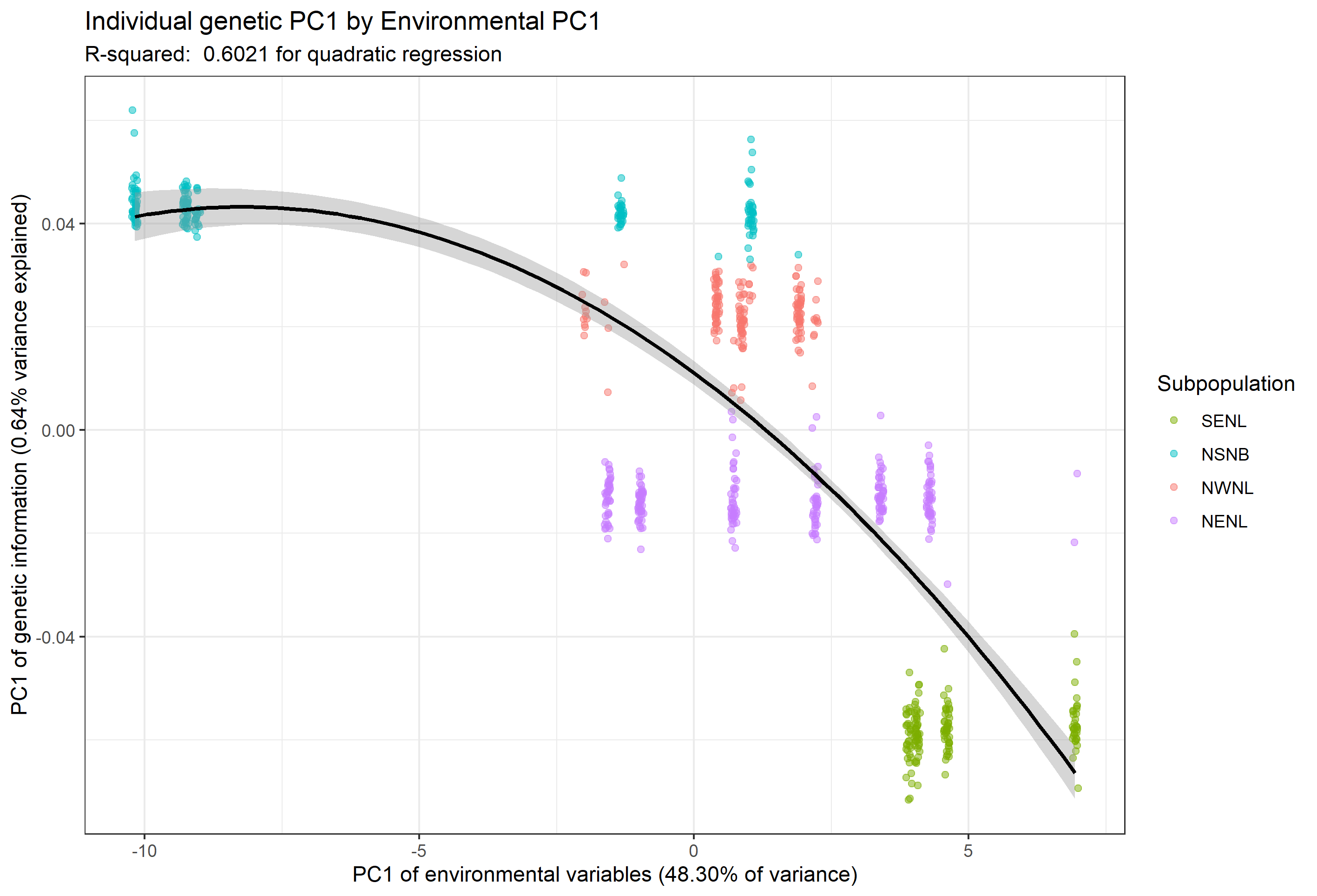

**Figure S10.** Scatter plot showing the relationship of the principal axes of environmental and genomic variation. Each point is an individual, and these are coloured by the subpopulation assignment for their location of origin (see Table 1). The x-axis is PC1 of environmental variation across the sampling locations (48.3% variance explained) and the y-axis is PC1 of genetic variation of all markers (n=11,574,435 markers) for all individuals (0.64% variance explained). The quadratic line of best fit is shown in black, with 95% confidence intervals in the flanking grey, the *r*^2^ of the correlation is 0.602 (p < 2.2e-16).

A)

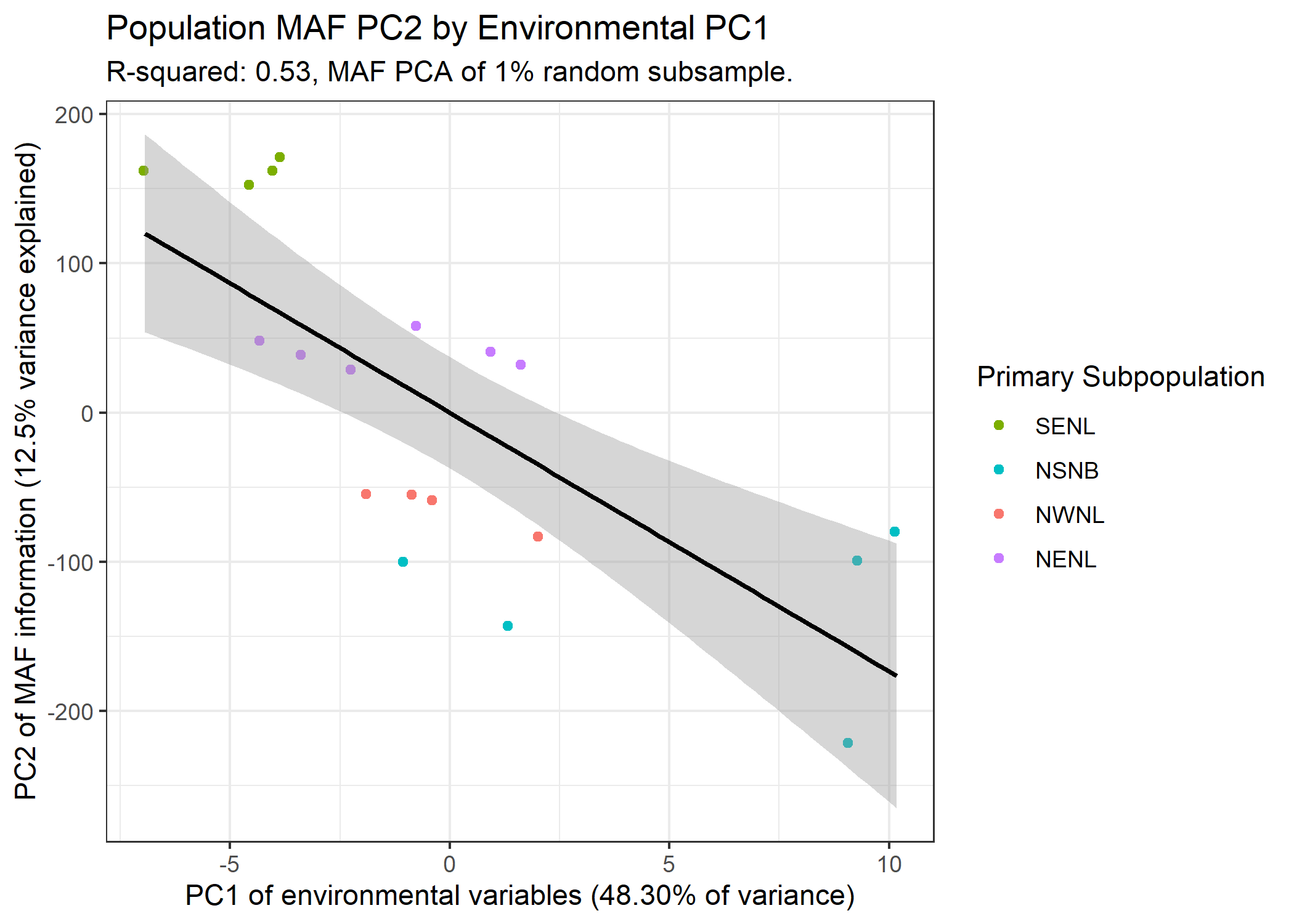

B)

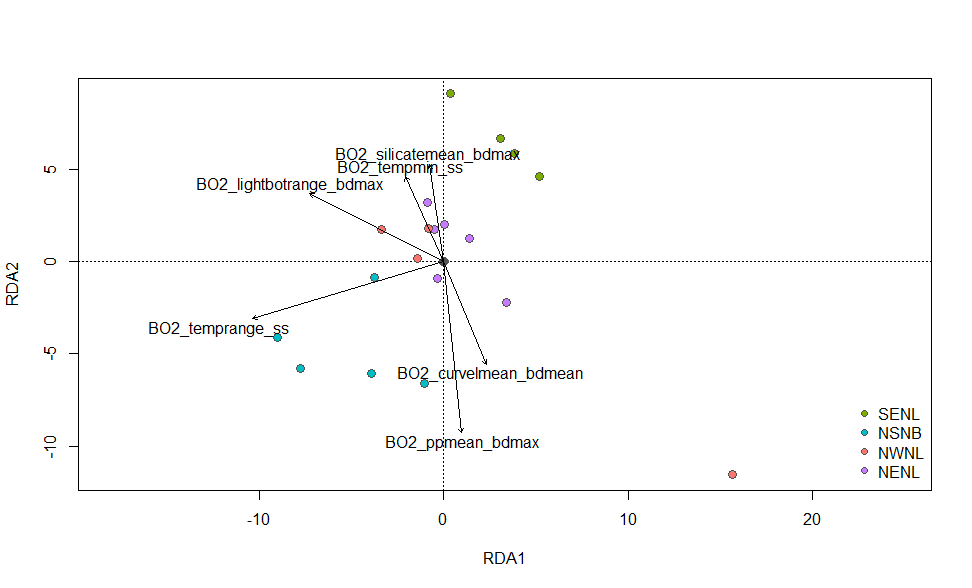

**Figure S11.** A) Scatter plot showing the relationship of the principal axes of environmental and per-location genomic variation. Each point is a location, and these are coloured by the primary subpopulation assignment for individuals from that location of origin (see Table 1). The x-axis is PC1 of environmental variation across the sampling locations (48.30% variance explained) and the y-axis is PC1 of the MAF variation amongst locations (12.5% variance explained). The linear line of best fit is shown in black, with 95% confidence intervals in the flanking grey. The *r*^2^ of the correlation is 0.53 (p < 2.2e-16). B) Ordination plot of the per-location (n=19) that used the six uncorrelated environmental variables (Table S3) as a predictor matrix and the MAF of a random 1% subsample of all markers as the response matrix. No correction for spatial structure was here applied. The adjusted *r*^2^ of the model was 0.045, suggesting the constrained ordination explained approximately 4.5% of the genetic variation. The small grey dots represent the minor allele frequencies, and the larger points represent the locations, coloured by their majority assigned population of origin. The black arrows represent the uncorrelated environmental predictors. The relative arrangement of the points and arrows on the plot represents their relationship with the RDA ordination axes (RDA1 = 0.295 proportion variance explained, RDA2 = 0.208 proportion variance explained).

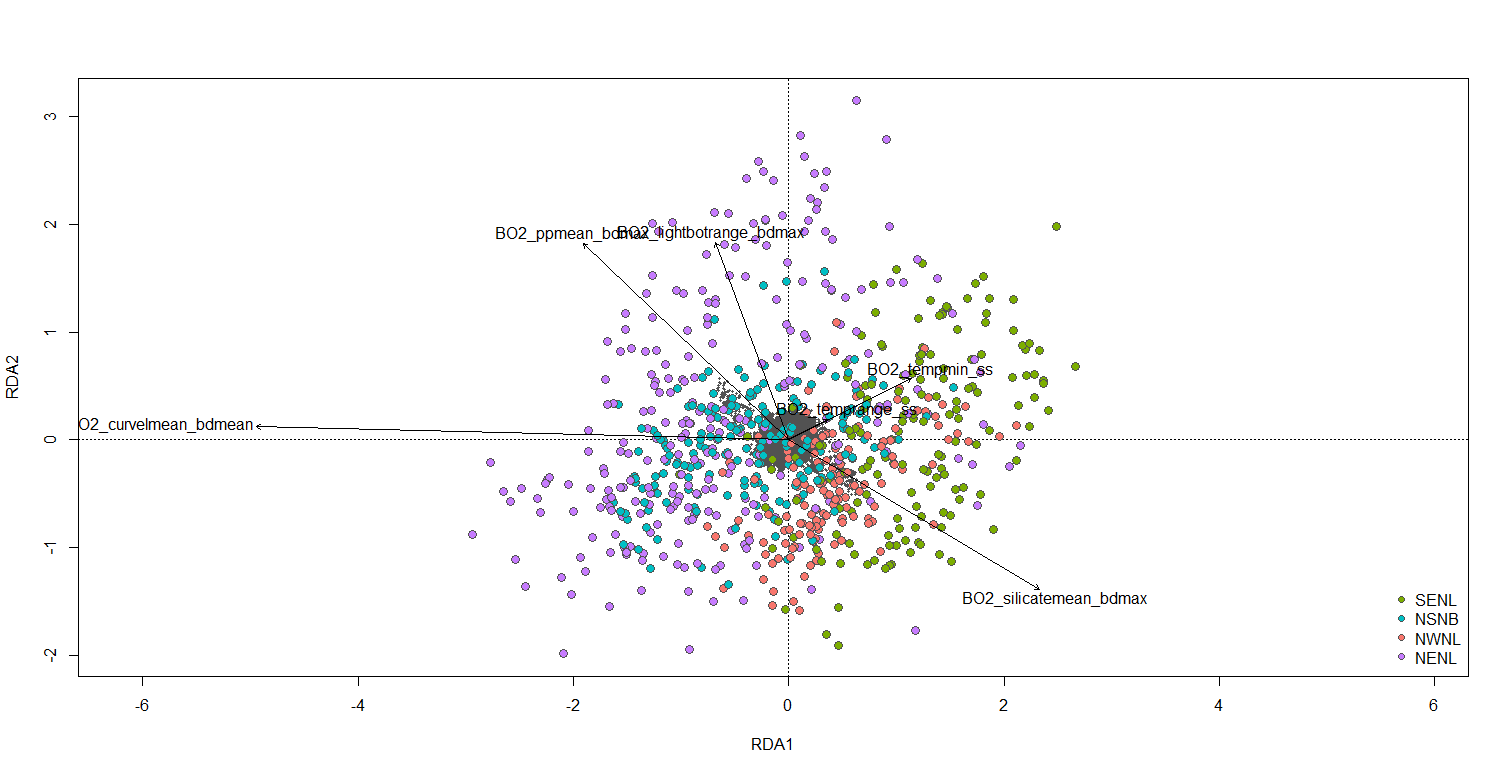
**Figure S12.** Ordination plot of the per-individual (n=749) RDA that used the six uncorrelated environmental variables (Table S3) as a predictor matrix and the genotype probabilities of the 7,698 markers from within the identified *F*_ST_ peaks as the response matrix. A MEM-based correction for spatial structure was here applied. The adjusted *r*^2^ of the model was 0.01, suggesting the constrained ordination explained approximately 1% of the genetic variation. Note that this value is likely an underestimate, as to increase resolution the response matrix was one-hot encoded (three columns per marker with the unique genotype probabilities) as opposed to dosage encoded. The small grey dots represent the genotype probabilities (response), and the larger points represent the individuals, coloured by their population of origin. The blue arrows represent the uncorrelated environmental predictors. The relative arrangement of the points and arrows on the plot represents their relationship with the RDA ordination axes (RDA1 = 0.5647 proportion variance explained, RDA2 = 0.1404 proportion variance explained).

**Table S3.** Correlation matrix of the different environmental variables from which a series of uncorrelated variables were selected for use within the rda. The vertical and horizontally aligned variable names indicate the pairwise correlation that corresponds to a given cell, cells coloured red indicate correlations more than 0.7 or less than -0.7. Asterisks indicate the set of 6 predictor variables retained for the RDA analysis.

| **curvelmean_bdmean*** | | | | | | | | | | | | | | | |  |  | | |  | | | | | | | | | |  | |  | | |  | | | |  | | | | | |  | | | |  | | | |  | | | |  | | | | |  | | | | | |  | | | | | |  | | | | | |  | | | | | |  | |  | | | | |  | | | | | |  | | | | | |  | | | | |  | | | | | |  | | | | | |  | | | | | |  | | | | |  | | | |
| --- | --- | --- | --- | --- | --- | --- | --- | --- | --- | --- | --- | --- | --- | --- | --- | --- | --- | --- | --- | --- | --- | --- | --- | --- | --- | --- | --- | --- | --- | --- | --- | --- | --- | --- | --- | --- | --- | --- | --- | --- | --- | --- | --- | --- | --- | --- | --- | --- | --- | --- | --- | --- | --- | --- | --- | --- | --- | --- | --- | --- | --- | --- | --- | --- | --- | --- | --- | --- | --- | --- | --- | --- | --- | --- | --- | --- | --- | --- | --- | --- | --- | --- | --- | --- | --- | --- | --- | --- | --- | --- | --- | --- | --- | --- | --- | --- | --- | --- | --- | --- | --- | --- | --- | --- | --- | --- | --- | --- | --- | --- | --- | --- | --- | --- | --- | --- | --- | --- | --- | --- | --- | --- | --- | --- | --- | --- | --- | --- | --- | --- | --- | --- | --- | --- | --- | --- |
| 0.07 | **dissoxmean_bdmax** | | | | | | | | | | | |  | | | |  | | | | | | | | | |  | | | |  | |  |  | | | |  |  | | |  | | | |  | | | |  | | | |  | | | |  | | | | | |  | | | | | |  | | | | | |  | | | | | |  | | | | | |  | | | | | |  | | | | | |  | | | | | |  | | | | | |  | | | | | |  | | | | | |  | | | |  | | | | |  | | | |
| 0.07 | 1.00 | | | **dissoxrange_bdmax** | | | | | | | | | | | | |  | | | | | | | | | |  | | | |  | |  |  | | | |  |  | | |  | | | |  | | | |  | | | |  | | | |  | | | | | |  | | | | | |  | | | | | |  | | | | | |  | | | | | |  | | | | | |  | | | | | |  | | | | | |  | | | | | |  | | | | | |  | | | | | |  | | | |  | | | | |  | | | |
| 0.24 | 0.91 | | | 0.91 | | | **dissoxltmax_bdmax** | | | | | | | | | | | | | | | | | | | |  | | | |  | |  |  | | | |  |  | | |  | | | |  | | | |  | | | |  | | | |  | | | | | |  | | | | | |  | | | | | |  | | | | | |  | | | | | |  | | | | | |  | | | | | |  | | | | | |  | | | | | |  | | | | | |  | | | | | |  | | | |  | | | | |  | | | |
| 0.14 | 0.53 | | | 0.53 | | | 0.49 | | | **ironmean_bdmax** | | | | | | | | | | | | | |  | | |  | | | | |  | | |  | | | |  | | |  | | | |  | | | |  | | | |  | | | |  | | | | | |  | | | | | |  | | | | | |  | | | | | |  | | | | | |  | | | | | |  | | | | | |  | | | | | |  | | | | | |  | | | | | |  | | | | | |  | | | |  | | | | |  | | | |
| 0.14 | 0.53 | | | 0.53 | | | 0.49 | | | 1.00 | | | **ironrange_bdmax** | | | | | | | | | | | | | |  | | | | |  | | |  | | | |  | | |  | | | |  | | | |  | | | |  | | | |  | | | | | |  | | | | | |  | | | | | |  | | | | | |  | | | | | |  | | | | | |  | | | | | |  | | | | | |  | | | | | |  | | | | | |  | | | | | |  | | | |  | | | | |  | | | |
| -0.41 | -0.68 | | | -0.68 | | | -0.63 | | | -0.54 | | | -0.54 | | | | **phosphatemean_bdmax** | | | | | | | | | | | | | | | | | |  | | | |  | | |  | | | |  | | | |  | | | |  | | | |  | | | | | |  | | | | | |  | | | | | |  | | | | | |  | | | | | |  | | | | | |  | | | | | |  | | | | | |  | | | | | |  | | | | | |  | | | | | |  | | | |  | | | | |  | | | |
| -0.41 | -0.68 | | | -0.68 | | | -0.63 | | | -0.54 | | | -0.54 | | | | 1.00 | | | | **phosphaterange_bdmax** | | | | | | | | | | | | | | | | | |  | | |  | | | |  | | | |  | | | |  | | | |  | | | | | |  | | | | | |  | | | | | |  | | | | | |  | | | | | |  | | | | | |  | | | | | |  | | | | | |  | | | | | |  | | | | | |  | | | | | |  | | | |  | | | | |  | | | |
| -0.08 | 0.14 | | | 0.14 | | | 0.08 | | | 0.10 | | | 0.10 | | | | -0.35 | | | | -0.35 | | | **lightbotmean_bdmax** | | | | | | | | | | | | | | |  | | |  | | | |  | | | |  | | | |  | | | |  | | | | | |  | | | | | |  | | | | | |  | | | | | |  | | | | | |  | | | | | |  | | | | | |  | | | | | |  | | | | | |  | | | | | |  | | | | | |  | | | |  | | | | |  | | | |
| -0.08 | 0.14 | | | 0.14 | | | 0.08 | | | 0.10 | | | 0.10 | | | | -0.35 | | | | -0.35 | | | 1.00 | | | **lightbotrange_bdmax*** | | | | | | | | | | | | | | |  | | | |  | | | |  | | | |  | | | |  | | | | | |  | | | | | |  | | | | | |  | | | | | |  | | | | | |  | | | | | |  | | | | | |  | | | | | |  | | | | | |  | | | | | |  | | | | | |  | | | |  | | | | |  | | | |
| -0.47 | -0.66 | | | -0.66 | | | -0.60 | | | -0.42 | | | -0.42 | | | | 0.89 | | | | 0.89 | | | -0.42 | | | -0.42 | | | | | **nitratemean_bdmax** | | | | | | | | | | | | | |  | | | |  | | | |  | | | |  | | | | | |  | | | | | |  | | | | | |  | | | | | |  | | | | | |  | | | | | |  | | | | | |  | | | | | |  | | | | | |  | | | | | |  | | | | | |  | | | |  | | | | |  | | | |
| -0.47 | -0.66 | | | -0.66 | | | -0.60 | | | -0.42 | | | -0.42 | | | | 0.89 | | | | 0.89 | | | -0.42 | | | -0.42 | | | | | 1.00 | | | **nitraterange_bdmax** | | | | | | | | | | | | | | |  | | | |  | | | |  | | | | | |  | | | | | |  | | | | | |  | | | | | |  | | | | | |  | | | | | |  | | | | | |  | | | | | |  | | | | | |  | | | | | |  | | | | | |  | | | |  | | | | |  | | | |
| -0.05 | 0.62 | | | 0.62 | | | 0.45 | | | 0.69 | | | 0.69 | | | | -0.56 | | | | -0.56 | | | 0.15 | | | 0.15 | | | | | -0.45 | | | -0.45 | | | | **tempmax_bdmax** | | | | | | | | | | | | | | |  | | | |  | | | | | |  | | | | | |  | | | | | |  | | | | | |  | | | | | |  | | | | | |  | | | | | |  | | | | | |  | | | | | |  | | | | | |  | | | | | |  | | | |  | | | | |  | | | |
| -0.05 | | 0.62 | | | 0.62 | | | 0.45 | | | 0.69 | | | 0.69 | | | | -0.56 | | | | -0.56 | | | 0.15 | | | 0.15 | | | | -0.45 | | | | -0.45 | | | | 1.00 | | | **tempmean_bdmax** | | | | | | | | | | | | | | | | | | | |  | | | | | |  | | | | | |  | | | | | |  | | | | | |  | | | | |  | | | | | |  | | | | | |  | | | | |  | | | | | |  | | | | | |  | | | | |  | | | | |  | | | |  | |
| -0.05 | | 0.62 | | | 0.62 | | | 0.45 | | | 0.69 | | | 0.69 | | | | -0.56 | | | | -0.56 | | | 0.15 | | | 0.15 | | | | -0.45 | | | | -0.45 | | | | 1.00 | | | 1.00 | | | | **tempmin_bdmax** | | | | | | | | | | | | | | | |  | | | | | |  | | | | | |  | | | | | |  | | | | | |  | | | | |  | | | | | |  | | | | | |  | | | | |  | | | | | |  | | | | | |  | | | | |  | | | | |  | | | |  | |
| -0.05 | | 0.62 | | | 0.62 | | | 0.45 | | | 0.69 | | | 0.69 | | | | -0.56 | | | | -0.56 | | | 0.15 | | | 0.15 | | | | -0.45 | | | | -0.45 | | | | 1.00 | | | 1.00 | | | | 1.00 | | | | **temprange_bdmax** | | | | | | | | | | | | | | | | | | | | | | | |  | | | | | |  | | | | | |  | | | | |  | | | | | |  | | | | | |  | | | | |  | | | | | |  | | | | | |  | | | | |  | | | | |  | | | |  | |
| 0.36 | | 0.58 | | | 0.58 | | | 0.59 | | | 0.60 | | | 0.60 | | | | -0.80 | | | | -0.80 | | | 0.15 | | | 0.15 | | | | -0.72 | | | | -0.72 | | | | 0.76 | | | 0.76 | | | | 0.76 | | | | 0.76 | | | | **carbonphytomean_bdmax** | | | | | | | | | | | | | | | | | | | | | | | | | | | |  | | | | | |  | | | | | |  | | | | | |  | | | | |  | | | | |  | | | | | |  | | | | | |  | | | | | |  | | | | |  | | |
| 0.36 | | 0.58 | | | 0.58 | | | 0.59 | | | 0.60 | | | 0.60 | | | | -0.80 | | | | -0.80 | | | 0.15 | | | 0.15 | | | | -0.72 | | | | -0.72 | | | | 0.76 | | | 0.76 | | | | 0.76 | | | | 0.76 | | | | 1.00 | | | | **carbonphytorange_bdmax** | | | | | | | | | | | | | | | | | | | | | | | | | | | | | |  | | | | | |  | | | | | |  | | | | |  | | | | |  | | | | | |  | | | | | |  | | | | | |  | | | | |  | | |
| 0.29 | | 0.60 | | | 0.60 | | | 0.58 | | | 0.76 | | | 0.76 | | | | -0.77 | | | | -0.77 | | | 0.12 | | | 0.12 | | | | -0.64 | | | | -0.64 | | | | 0.82 | | | 0.82 | | | | 0.82 | | | | 0.82 | | | | 0.96 | | | | | 0.96 | | | | | | **ppmean_bdmax*** | | | | | | | | | | | | | | | | | |  | | | | | |  | | | | | |  | | | | | |  | | | | |  | | | | | |  | | | | | |  | | | | | |  | | | | |  | | | | |  | |
| 0.06 | | -0.82 | | | -0.82 | | | -0.67 | | | -0.48 | | | -0.48 | | | | 0.64 | | | | 0.64 | | | -0.37 | | | -0.37 | | | | 0.71 | | | | 0.71 | | | | -0.76 | | | -0.76 | | | | -0.76 | | | | -0.76 | | | | -0.64 | | | | -0.64 | | | | | | -0.64 | | | | | | **salinitymean_bdmax** | | | | | | | | | | | | | | | | | | | | | | | |  | | | | | |  | | | | |  | | | | |  | | | | | |  | | | | | |  | | | | | |  | | | | |  | | |
| -0.50 | | -0.22 | | | -0.22 | | | -0.27 | | | -0.37 | | | -0.37 | | | | 0.60 | | | | 0.60 | | | -0.07 | | | -0.07 | | | | 0.49 | | | | 0.49 | | | | -0.53 | | | -0.53 | | | | -0.53 | | | | -0.53 | | | | -0.70 | | | | -0.70 | | | | | | -0.62 | | | | | | 0.30 | | | | | | **silicatemean_bdmax*** | | | | | | | | | | | | | | | | | | | | | | | |  | | | | |  | | | | |  | | | | | |  | | | | | |  | | | | | |  | | | | |  | | |
| -0.19 | | 0.57 | | | 0.57 | | | 0.57 | | | 0.19 | | | 0.19 | | | | -0.03 | | | | -0.03 | | | 0.02 | | | 0.02 | | | | -0.21 | | | | -0.21 | | | | 0.43 | | | 0.43 | | | | 0.43 | | | | 0.43 | | | | 0.32 | | | | | 0.32 | | | | | | 0.30 | | | | | | -0.62 | | | | | | 0.05 | | | | | | **icecovermax_ss** | | | | | | | | | | | | | | | | | |  | | | | |  | | | | | |  | | | | | |  | | | | | |  | | | | |  | | | | |  | |
| -0.05 | | 0.80 | | | 0.80 | | | 0.76 | | | 0.42 | | | 0.42 | | | | -0.50 | | | | -0.50 | | | -0.05 | | | -0.05 | | | | -0.45 | | | | -0.45 | | | | 0.70 | | | 0.70 | | | | 0.70 | | | | 0.70 | | | | 0.66 | | | | | 0.66 | | | | | | 0.67 | | | | | | -0.76 | | | | | | -0.29 | | | | | | 0.73 | | | | | | **icecovermean_ss** | | | | | | | | | | | | | | | | |  | | | | | |  | | | | | |  | | | | | |  | | | | |  | | | | |  | |
| -0.19 | | 0.57 | | | 0.57 | | | 0.57 | | | 0.19 | | | 0.19 | | | | -0.03 | | | | -0.03 | | | 0.02 | | | 0.02 | | | | -0.21 | | | | -0.21 | | | | 0.43 | | | 0.43 | | | | 0.43 | | | | 0.43 | | | | 0.32 | | | | | 0.32 | | | | | | 0.30 | | | | | | -0.62 | | | | | | 0.05 | | | | | | 1.00 | | | | | | 0.73 | | | | | | **icecoverrange_ss** | | | | | | | | | | | | | | | | |  | | | | | |  | | | | | |  | | | | |  | | | | |  | |
| 0.27 | | 0.79 | | | 0.79 | | | 0.92 | | | 0.33 | | | 0.33 | | | | -0.52 | | | | -0.52 | | | 0.06 | | | 0.06 | | | | -0.57 | | | | -0.57 | | | | 0.38 | | | 0.38 | | | | 0.38 | | | | 0.38 | | | | 0.59 | | | | | 0.59 | | | | | | 0.55 | | | | | | -0.64 | | | | | | -0.25 | | | | | | 0.72 | | | | | | 0.80 | | | | | | 0.72 | | | | | | **icethickmax_ss** | | | | | | | | | | | | | | | | |  | | | | | |  | | | | |  | | | | |  | |
| 0.16 | | 0.71 | | | 0.71 | | | 0.78 | | | 0.27 | | | 0.27 | | | | -0.57 | | | | -0.57 | | | 0.00 | | | 0.00 | | | | -0.53 | | | | -0.53 | | | | 0.48 | | | 0.48 | | | | 0.48 | | | | 0.48 | | | | 0.67 | | | | | 0.67 | | | | | | 0.63 | | | | | | -0.65 | | | | | | -0.36 | | | | | | 0.59 | | | | | | 0.88 | | | | | | 0.59 | | | | | | 0.89 | | | | | **icethickmean_ss** | | | | | | | | | | | | | | | | | |  | | | | |  | | | | |  | |
| 0.27 | | 0.79 | | | 0.79 | | | 0.92 | | | 0.33 | | | 0.33 | | | | -0.52 | | | | -0.52 | | | 0.06 | | | 0.06 | | | | -0.57 | | | | -0.57 | | | | 0.38 | | | 0.38 | | | | 0.38 | | | | 0.38 | | | | 0.59 | | | | | 0.59 | | | | | | 0.55 | | | | | | -0.64 | | | | | | -0.25 | | | | | | 0.72 | | | | | | 0.80 | | | | | | 0.72 | | | | | | 1.00 | | | | | 0.89 | | | | | | **icethickrange_ss** | | | | | | | | | | | | | | | | |  | | | | |  | |
| -0.51 | | 0.43 | | | 0.43 | | | 0.18 | | | 0.25 | | | 0.25 | | | | 0.04 | | | | 0.04 | | | 0.19 | | | 0.19 | | | | 0.03 | | | | 0.03 | | | | 0.57 | | | 0.57 | | | | 0.57 | | | | 0.57 | | | | 0.01 | | | | | 0.01 | | | | | | 0.08 | | | | | | -0.54 | | | | | | 0.16 | | | | | | 0.41 | | | | | | 0.32 | | | | | | 0.41 | | | | | | 0.01 | | | | | -0.04 | | | | | | 0.01 | | | | | | **tempmax_ss** | | | | | | | | | | | | | | | |  | |
| -0.50 | | | 0.27 | | | 0.27 | | | -0.04 | | | 0.27 | | | 0.27 | | | | 0.04 | | | | 0.04 | | | 0.15 | | | 0.15 | | | 0.08 | | | | | 0.08 | | | | 0.53 | | | 0.53 | | | | 0.53 | | | | 0.53 | | | | -0.06 | | | | | -0.06 | | | | | | 0.05 | | | | | | -0.38 | | | | | | 0.13 | | | | | | 0.14 | | | | | | 0.11 | | | | | | 0.14 | | | | | | -0.26 | | | | | -0.27 | | | | | | -0.26 | | | | | | 0.93 | | | | | | | **tempmean_ss** | | | | | | | | | |
| -0.23 | | | -0.53 | | | -0.53 | | | -0.76 | | | -0.35 | | | -0.35 | | | | 0.48 | | | | 0.48 | | | -0.09 | | | -0.09 | | | 0.47 | | | | | 0.47 | | | | -0.04 | | | -0.04 | | | | -0.04 | | | | -0.04 | | | | -0.41 | | | | | -0.41 | | | | | | -0.38 | | | | | | 0.33 | | | | | | 0.13 | | | | | | -0.36 | | | | | | -0.48 | | | | | | -0.36 | | | | | | -0.78 | | | | | -0.63 | | | | | | -0.78 | | | | | | 0.29 | | | | | | | 0.49 | | | | | **tempmin_ss*** | | | | |
| -0.49 | | | 0.56 | | | 0.56 | | | 0.34 | | | 0.33 | | | 0.33 | | | | -0.06 | | | | -0.06 | | | 0.21 | | | 0.21 | | | -0.07 | | | | | -0.07 | | | | 0.60 | | | 0.60 | | | | 0.60 | | | | 0.60 | | | | 0.09 | | | | | 0.09 | | | | | | 0.16 | | | | | | -0.62 | | | | | | 0.14 | | | | | | 0.50 | | | | | | 0.42 | | | | | | 0.50 | | | | | | 0.17 | | | | | 0.09 | | | | | | 0.17 | | | | | | 0.98 | | | | | | | 0.87 | | | | | 0.10 | | | | **temprange_ss*** |
